# Disruption of the transcription factor *NEUROD2* causes an autism syndrome via cell-autonomous defects in cortical projection neurons

**DOI:** 10.1101/296889

**Authors:** Karen Runge, Rémi Mathieu, Stéphane Bugeon, Sahra Lafi, Corinne Beurrier, Surajit Sahu, Fabienne Schaller, Arthur Loubat, Leonard Herault, Stéphane Gaillard, Mélanie Cahuc, Emilie Pallesi-Pocachard, Aurélie Montheil, Andreas Bosio, Jill A Rosenfeld, Eva Hudson, Kristin Lindstrom, Saadet Mercimek-Andrews, Lauren Jeffries, Arie van Haeringen, Olivier Vanakker, Bruno Pichon, Audrey Van Hecke, Dina Amrom, Sebastien Küry, Candace Gamble, Bernard Jacq, Laurent Fasano, Gabriel Santpere, Belen Lorente-Galdos, Nenad Sestan, Antoinette Gelot, Sylvie Giacuzzo, Alfonso Represa, Carlos Cardoso, Harold Cremer, Antoine de Chevigny

## Abstract

We identified seven families associating *NEUROD2* pathogenic mutations with ASD and intellectual disability. To get insight into the pathophysiological mechanisms, we analyzed cortical development in *Neurod2* KO mice. Cortical projection neurons (CPNs) over-migrated during embryogenesis, inducing abnormal thickness and laminar positioning of cortical layers. At juvenile ages, dendritic spine turnover and intrinsic excitability were increased in L5 CPNs. Differentially expressed genes in *Neurod2* KO mice were enriched for voltage-gated ion channels, and the human orthologs of these genes were strongly associated with ASD. Furthermore, adult *Neurod2* KO mice exhibited core ASD-like behavioral abnormalities. Finally, by generating *Neurod2* conditional mutant mice we demonstrate that forebrain excitatory neuron-specific *Neurod2* deletion recapitulates cellular and behavioral ASD phenotypes found in full KO mice. Our findings demonstrate crucial roles for *Neurod2* in cortical development and function, whose alterations likely account for ASD and related symptoms in the newly defined *NEUROD2* mutation syndrome.

## Introduction

Alterations in cellular development, synaptic transmission and intrinsic excitability of cortical projection neurons (CPNs) (*1, 2*) are prevalent theories of the pathophysiology of neurodevelopmental disorders including autism spectrum disorders (ASD) and intellectual disability (*3–5*). Despite a field of intense investigation, the transcription factors that regulate these processes remain poorly known.

NEUROD2 belongs to the family of NEUROD basic helix-loop-helix transcription factors that regulate early neuronal differentiation during development (*6*). While cortical expression of its closest and first identified paralog NEUROD1 is turned off around birth (*6*), NEUROD2 cortical expression persists postnatally (*7*), indicating that it might be involved in other processes. Few studies have suggested a link between NEUROD2 and synapse formation. Indeed, *Neurod2* mutant mice have a reduced density of synaptic markers in amygdala (*8*), and the degradation of NEUROD2 by the ubiquitin-proteasome system is required for the maturation of presynaptic terminals in cerebellar neurons (*9*). Also, the electrophysiological maturation of the thalamo-cortical synapse is altered in the barrel cortex of *Neurod2* deficient mice (*10*).

A possible link between *NEUROD2* and neuropsychiatric disorders has recently emerged. In a recent study two children with early epileptic encephalopathy were found to carry *de novo NEUROD2* variants, and functional analyses in tadpoles suggested that these variants were pathogenic (*11*). Moreover, several lines of evidence suggest that *NEUROD2* might be associated with schizophrenia, intellectual disability and/or ASD. First, four rare *NEUROD2* polymorphisms have been associated with schizophrenia (*12*). Second, analysis of miRNA prediction algorithms (TargetScan, miRanda) shows that *NEUROD2* mRNA contains a high-confidence putative target site for the schizophrenia-related microRNA miR-137 (*13*). Third, the main transcriptional cofactor of NEUROD2 is the Pitt-Hopkins syndrome gene TCF4 that confers intellectual disability (*14*). Fourth, copy number variations encompassing *NEUROD2* are associated with intellectual disability and/or ASD in the DECIPHER database (https://decipher.sanger.ac.uk/). Fifth, a machine learning based algorithm for the prediction of ASD-related genes ranked *NEUROD2* 98^th^ among all human genes (*15*) – a better rank than many confirmed ASD genes -, which positions *NEUROD2* in the core of genetic networks and signaling pathways associated with ASD. Together, these pieces of evidence support the hypothesis that *NEUROD2* might be associated with neuropsychiatric disorders in humans.

Besides hippocampus and amygdala, the cerebral cortex is the brain structure that is the most strongly associated with neuropsychiatric diseases (*1, 2*). However, current knowledge about the role of *Neurod2* in cortex is limited. *Neurod2* deficient mice have a disorganized barrel cortex although production of CPNs in layer 4 (L4) is normal (*10*). Electrophysiologically, L2/3 CPNs show increased intrinsic excitability in somatosensory cortex (*16*). In spite of this information, the impact of *Neurod2* deletion on cortical development has not been investigated.

Here we identified seven families with pathogenic *NEUROD2* mutations causing as core clinical symptoms ASD, intellectual disability and speech delay. By combining mouse genetics, RNA-seq analyses, electrophysiology and behavioral testing, we provide strong experimental evidence for a causal relationship between *Neurod2* disruption, functional defects in CPNs and ASD.

## Material, Patients & Methods

### Consent and human ethics approval

All subjects or their legal representatives gave written informed consent for the study. Except for family 3 and 4 for which the study was approved by the institutional review Board (IRB) at Yale University, studies of other families in the present work used unlinked anonymized data and were performed in accordance with the Declaration of Helsinki protocols and approved by the Dutch, USA and Belgian ethics committees. The clinical cytogenetic sample consisted of patients referred to the Netherlands, USA, and Belgium from regional pediatricians, other health specialists and/or genetics centers.

### Patients

The study includes 9 patients from 6 unrelated families in the Netherlands, USA, Mexico, Canada and Belgium. These patients were identified in centers where High-throughput Sequencing (whole-exome sequencing, whole-genome sequencing) is being used to identify, or accurately characterize, genomic variants in individuals with developmental brain abnormalities, mental retardation, epilepsy and ASD. Clinical information, brain MRI, and blood were obtained after informed consent. DNA from subjects was extracted from peripheral blood lymphocytes by standard extraction procedures.

### Animals

Mice (mus musculus) were group housed (2–5 mice/cage) with same-sex littermates on a 12 hour light-dark cycle with access to food and water ad libitum. *Neurod2* full KO mice were previously described (*17*). They were bred and maintained on a mixed SVeV-129/C57BL/6J background, which allows mouse survival (*18*). *Neurod2*^flox/flox^ mice were generated using cell clone EPD0422fl5flB06 from the Knock Out Mouse Project at UC Davis (http://www.komp.org), in which the only coding exon (exon 2) was flanked by loxP sites (Supplementary Fig. 8). Experimenters were always blinded to genotypes during data acquisition and analysis. Animal experiments were carried out in accordance with European Communities Council Directives and approved by French ethical committees (Comité d’Ethique pour l’expérimentation animale no. 14; permission number: 62-12112012). Male mice were used for behavioral experiments, while mice of either sex were used for all other experiments.

### Immunostainings

Mice were perfused transcardially with ice-cold 4% paraformaldehyde (in PBS). Brains were removed and post-fixed overnight at 4°C with the same fixative. Coronal sections were cut at 50 μm thickness using a cryostat (Leica) or a sliding microtome (Microm).

Immunofluorescence experiments were performed as described before (*19*). Briefly, free-floating sections were blocked and permeabilized for one hour in a blocking buffer composed of 10% Normal Goat Serum, 0.2% Triton X-100 (Sigma) in PBS. Primary antibodies, diluted in blocking solution and added overnight at 4°C, were as follows: rabbit anti-NEUROD2 (1:500, Abcam, #ab104430), rabbit anti-TBR1 (1:1000, Abcam, #31940), rat anti-BCL11B (1:100, Abcam, #ab18465), rabbit anti-CUX1 (1:200, Santa Cruz Biotechnology, #sc13024), mouse anti-RORβ (1:200, Perseus Proteomics, #PP-N7927-00), chicken anti-GFP (1:500, Aves, #GFP-1010). Corresponding fluorescently labeled secondary antibodies (AlexaFluor, Invitrogen) were added for 2 hours in blocking solution. Hoechst was added in PBS for 10 minutes, and sections were mounted on microscope slides that were coversliped using Mowiol solution (Sigma).

### *In Utero* Electroporation

Timed pregnant *Neurod2* heterozygous females fertilized by heterozygous males (E13.5) were anesthetized with isoflurane (7.5% for induction and 3.5% for surgery). The uterine horns were exposed. A volume of 1–2 µL of DNA plasmid (pCAGGS-RFP, 0.5 μg/μL) combined with Fast Green (2 mg/ mL, Sigma) was injected into the lateral ventricle of each embryo with a pulled glass capillary and a microinjector (Picospritzer II, General Valve Corporation, Fairfield, NJ, USA). Electroporation was then conducted by discharging a 4000 µF capacitor charged to 50 V with a BTX ECM 830 electroporator (BTX Harvard Apparatus, Holliston, MA, USA). Five electric pulses (5 ms duration) were delivered at 950 ms intervals using electrodes. Embryos were collected 5 days post-electroporation (at E18.5), and 80 μm coronal slices cut using a sliding microtome (Microm). Electroporated cortices were imaged with an Apotome microscope (Zeiss). Quantification of red fluorescent protein (RFP)-expressing cell distribution was performed on cortical column crop images (bottom: above VZ/SVZ, below IZ; top: brain surface) using a custom-made algorithm on Fiji. We counted the proportion of total RFP-positive cells that were found in each of 20 bins of equal size along the width of the cortical column.

### Laminar positioning, radial migration and layer thickness measurements

Images used to analyze the lamination of CPN subtypes at P30 and of *in utero* electroporated neurons at E18.5 were acquired using an apotome microscope (Zeiss) with a 10x objective. Typically, three stacks with 1 μm z-steps centered in the mid depth of each slice were imaged and a maximum intensity projection was generated. Cortical columns were cropped in stereotyped positions of the somatosensory cortex. For P30 mice lower and upper limits of the cropped images were the dorsal border of corpus callosum and the L1-L2 boundary, respectively. For E18.5 pups, crop limits were SVZ-IZ boundary and pial surface, respectively. Blind manual counting of marker-positive cells was made in 20 bins using Image J.

For radial cell quantification of PNs, 5 slices per animal were selected in defined levels of the rostro-caudal axis. For each slice, a cortical column was cropped in the somatosensory cortex by using PhotoShop software. Each cortical column was divided into 20 Bins of equal size spanning the cortical thickness. Cells were quantified in each bin, and bin-distribution was defined as the percentage of cells in each bin relative to the total number of cells.

For layer thickness measurement, the same slices were used. For each slice, the layer thickness was measured at the left edge, the right edge and the middle of the cortical column and the mean of these 3 values was calculated. Limits of the layer thickness were defined by only taking into account strong fluorescent cells. Layer thickness of each slice was then calculated relatively to the whole cortex thickness. For radial cell quantification of electroporated cells, slices were selected at 3 defined levels of the rostro-caudal axis. The same method than the one for radial cell quantification of PNs was applied as previously detailed.

All counts were manually performed by an investigator who was blinded to genotype.

### Dendritic reconstructions and length measurement

For 3D dendritic reconstructions, Thy1-GFP expressing PNs were imaged as 3D stacks using a Zeiss LSM 800 confocal microscope with a 20X objective and a 0.5 μm z-step. Neurons were reconstructed three-dimensionally using Imaris software (Bitplane). Digital reconstructions were analyzed with Imaris to measure the total dendritic length.

### Spine density measurements

For spine analyses, images were acquired as 3D stacks with lateral and z-axis resolution of 100 nm and 380 nm using a Zeiss LSM 800 confocal microscope, with an oil-immersed 63X objective. For the quantitative and morphological analyses of dendritic tree spines, basal and apical dendrites were imaged separately using a 63X oil objective, roughly placing the soma at the center of the image. During the same imaging session, randomly selected dendrites were imaged by using the same objective and adding a 3.4 numerical zoom. For basal dendritic spines, dendrites 5–100 μm distal to the first bifurcation away from the soma were imaged. For apical dendritic spines, dendrites within 5-100 μm of the pial dendritic ending were imaged. Spine quantifications were performed on z-stacks using NeuronStudio. Imaging and quantifications were all performed blindly to experimental groups.

### In vivo trans-cranial two-photon imaging for spine turnover measurements

Four-week-old *Neurod2 KO*; Thy1-GFP mice were ketamine/xylazine anesthetized. The skin over the skull was incised, and the skull was cleaned to ensure adhesion, using activator (10% citric acid). An aluminum bar was then glued and cemented using dental cement (Super-Bond C&B) on the skull, caudally to Bregma, keeping the skull parallel to the bar. The animal was then placed in a stereotaxic frame, attached to the metal bar, and the primary motor cortex was marked (coordinates: +1.0 AP; +2.0 ML). Using a scalpel blade, the skull was gently thinned until less than 20 μm thickness, avoiding any bleeding, breaking or extensive drying of the skull. Dendritic segments that were bright and visible were located and marked on a low magnification z-stack and then imaged at high resolution (axial resolution: 0.21 μm; step size: 0.79 μm). After imaging the skin over the skull was stitched and animals received a subcutaneous injection of Carprofen (5 mg/kg) and Buprenorphine (0.3 mg/kg). Three days after the first imaging session, animals were anesthetized and the skin covering the skull was re-opened. Next, the skull was cleaned and new images of each dendritic segment were taken. New and eliminated spines were then determined blindly, comparing images at day 0 and at day 3 on ImageJ, on single z-planes.

### Image Analyses

All cells (lamination and radial migration experiments), dendrites and spines analyzed were located in the M1 and S1 cortex. Image analyses were always performed with the scientists blinded to the experimental conditions (we used a custom-made algorithm to blind all data). Manual quantifications of CPN lamination, dendrite and spine parameters by 2 independent qualified scientists gave similar results.

### Quantitative reverse transcription polymerase chain reaction (qRT-PCR)

Total RNAs were extracted from whole cortices (for developmental expression of *Neurod2* in Fig. S2a, n = 3-4 brains per stage in each condition) or from the pooled motor and somatosensory areas (for confirmation of RNAseq data in Fig. 4e, n=3 samples) using TRIZOL reagent according to manufacturer’s instructions (Life Technology). cDNA was synthesized from 1 μg of total RNA using Quantitect Reverse Transcription Kit and according to manufacturer protocol (Qiagen). RT-PCRs were then carried out using SYBR-Green chemistry (Roche Diagnostics) and Roche amplification technology (Light Cycler 480). PCR primers (Table below) were designed for 12 mouse genes, and for 3 control genes, *Ptgs2* (Prostaglandin-Endoperoxide Synthase 2), Rpl13a (Ribosomal protein L13a) and *Hprt1* (Hypoxanthine Phosphoribosyltransferase 1) for normalization. All primer pairs were optimized to ensure specific amplification of the PCR product and the absence of any primer dimer. Quantitative PCR standard curves were set up for all. Values of fold change represent averages from duplicate measurements for each sample.

**Table 1:**
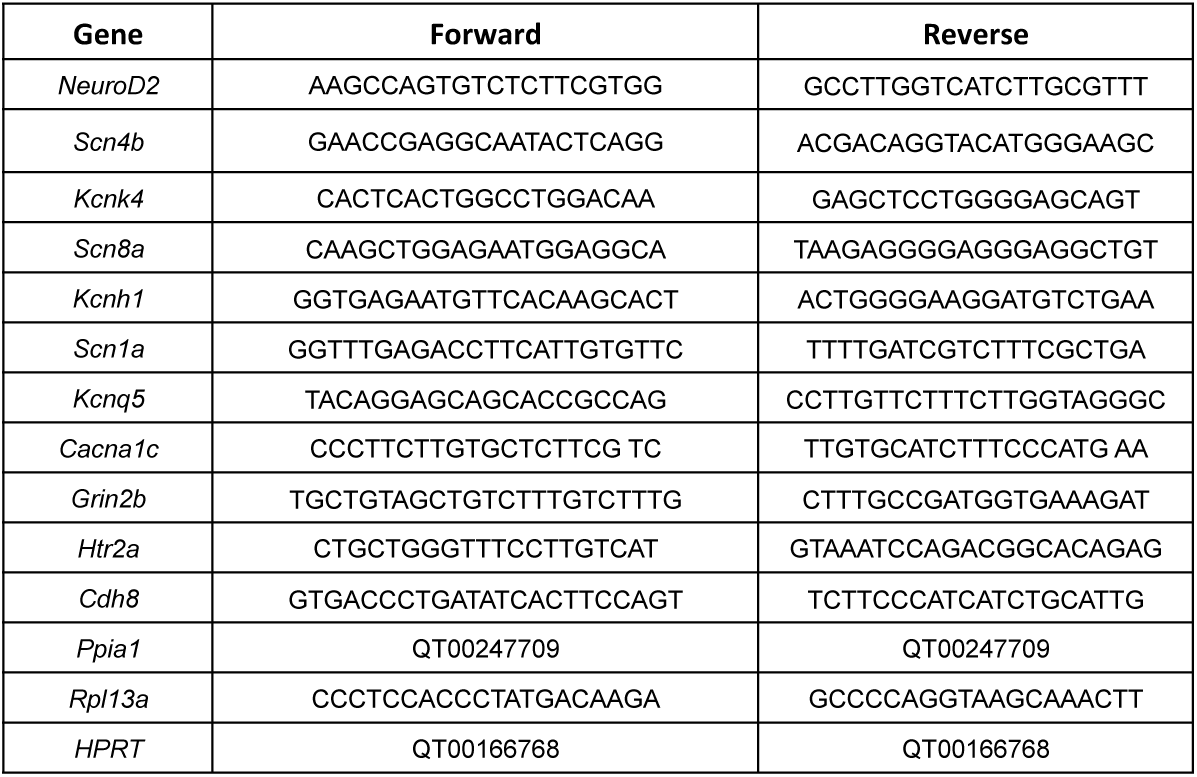
Primer pairs used for qPCR in this study.

#### Electrophysiological recordings

Coronal slices (250 µm) from 21 to 30 days-old mice were cut with a VT 1000S vibratome (Leica) in ice-cold high-choline artificial cerebro-spinal fluid (ACSF) containing (in mM): 130 choline, 2.5 KCl, 1.25 NaH2PO4, 7 MgCl2, 0.5 CaCl2, 25 NaHCO3 and 7 glucose at 4°C. Slices were then maintained at room temperature in oxygenated ACSF containing (in mM): 126 NaCl, 2.5 KCl, 1.2 NaH2PO4, 1.2 MgCl2, 2.4 CaCl2, 25 NaHCO3 and 11 glucose, to which 250 µM kynurenic acid and 1 mM sodium pyruvate were added. Slices were then transferred one at a time to a submersion recording chamber and were perfused continuously with ACSF warmed to 33°C at a rate of 2.5-3 ml/min. All solutions were equilibrated with 95% O2/ 5% CO2. Neurons were visualized on an upright microscope (Nikon Eclipse FN1) equipped with DIC optic and filter set to visualize EYFP using a x40 water-immersion objective. Recordings were interleaved in control and *Neurod2* KO mice.

Miniature excitatory and inhibitory postsynaptic currents (mEPSCs and mIPSCs, respectively) were recorded in whole-cell configurations in voltage-clamp mode in oxygenated ACSF containing tetrodotoxin (TTX, 1 µM). Patch-clamp electrodes (4-6 MΩ) were filled with an intracellular solution of the following composition (in mM): 120 CsMeSO4, 12.3 CsCl, 0.1 CaCl2, 1 EGTA, 10 HEPES, 4 MgATP, 0.3 NaGTP, pH adjusted to 7.25 with CsOH and osmolarity adjusted to 270-280 mOsm/L. Cells were kept at −60 mV, the reversal potential for GABAergic events, or −4 mV, the reversal potential for glutamatergic events, for the recordings of mEPSCs and mIPSCs, respectively. In some experiments, picrotoxin (50 µM) and 6-cyano-7-nitroquinoxaline-2,3-dione (CNQX, 10 µM) were applied at the end of the experiment to verify that the currents were indeed GABAergic and glutamatergic, respectively. Access resistance was monitored throughout the experiments with a 5-mV negative step and was found to be constant.

To measure intrinsic properties, we used current-clamp recordings. Glass electrodes (6–9 MΩ) were filled with an internal solution containing the following (mM): 130 KMeSO4, 5 KCl, 10 4-(2-hydroxyethyl)-1-piperazineethanesulfonic acid, 2.5 MgATP, 0.3 NaGTP, 0.2 ethyleneglycoltetraacetic acid, 10 phosphocreatine, and 0.3-0.5% biocytin, pH = 7.21. Access resistance ranged between 15-22 MΩ, and the results were discarded if the access resistance changed by >20%. TTX, was obtained from Abcam, CNQX from Tocris and picrotoxin and kynurenic acid from Sigma.

Data were collected with a MultiClamp 700B amplifier (Molecular Devices), filtered at 2kHz, digitized (10kHz) with a Digidata 1440A (Molecular Devices) to a personal computer, and acquired using Clampex 10.1 software (PClamp, Axon Instruments, Molecular Devices). Data were analyzed and plotted in clampfit (Molecular Devices, v 10.2). Miniature currents were analyzed with Mini Analysis (Synaptosoft, version 6.0.7).

### Retrograde tracing

P28 mice under xylazine/ketamine anesthesia received stereotaxic injections of 0.3 µl of cholera toxin subunit B (CT-B, 1 mg/ml; Thermo Fisher Scientific) conjugated with Alexa Fluor 647 in the striatum (AP: +1 mm; ML: +1.8 mm; DV: −2.9 mm from dura) and conjugated with Alexa Fluor 488 in the thalamus (AP: −1.3 mm; ML: +1.15 mm; DV: −3.5 mm from dura) using Bregma coordinates. This allowed retrograde labeling of, respectively, L5 PNs (striatal injection) and L6 PNs (thalamic injection). Another group of animals were injected with Alexa Fluor 488 CT-B in motor cortex (AP: 0.6 mm; ML: 1.3 mm; DV: 0.7 mm). Ten days after injection, animals were trans-cardially perfused with 4% paraformaldehyde, their brains processed and cut 50 μm thick using a sliding microtome (Microm).

### RNA Isolation and library preparation

Tissue from motor plus somatosensory cortex of P28 mice was rapidly micro-dissected and frozen at −80°C (n=3 independent experiments for WT and KO samples, 1 to 4 mice per sample). Total RNA was purified using spin columns of the RNeasy Mini Kit (Qiagen) according to manufacturer’s protocol. Library preparation was made with the TruSeq mRNA-seq Stranded v2 Kit sample preparation (Illumina) according to manufacturer’s instructions. One µg total RNA was used for poly(A)-selection and Elution-Fragmentation incubation time was 8 min to obtain 120-210 bp fragments. Each library was barcoded using TruSeq Single Index (Illumina). After library preparation, Agencourt AMPure XP (Beckman Coulter, Inc.) was performed for 200 to 400 bp libraries size-selection (282 nt average final library size). Each library was examined on the Bioanalyzer with High Sensitivity DNA chip (Agilent), quantified on Qubit with Qubit® dsDNA HS Assay Kit (Life Technologies), diluted to 4 nM and then pulled together at equimolar ratio.

### Illumina NextSeq-500 sequencing

Sequencing was performed by the TGML Facility (INSERM U1090) using PolyA mRNA isolation, directional RNA-seq library preparation and the Illumina NextSeq 500 sequencer. The denaturation was performed with 5 µl of pooled libraries (4 nM) and 5 min incubation with 5 µl of fresh NaOH (0.2N) and then addition of 5 µL of fresh Tris-HCl (200 mM - pH 7), according to manufacturer’s instructions. The dilution of 20 pM pooled libraries was performed with HT1 to a 1.2 pM final concentration. PhiX library as a 1% spike-in for use as a sequencing control was denatured and diluted, and 1.2 µl was added to denature and dilute pooled libraries before loading. Finally, libraries were sequenced on a high-output flow cell (400M clusters) using the NextSeq® 500/550 High Output v2 150 cycles kit (Illumina), in paired-end 75/ 75nt mode, according to manufacturer’s instructions.

### RNA-seq data primary analysis

467 548 934 clusters were generated for 71 Gbp sequenced with 75 % >= Q30. Reads were first trimmed with sickle v1.33 (Joshin et al., 2011) (RRintellectual disability:SCR_006800) with parameters –l 25 –q 20 and then aligned to mm10 using STAR v2.5.3a (Dobin et al., 2013) to produce BAM alignment files. Multi-mapped reads and reads with more than 0.08 mismatches per pair relative to read length were discarded. Transcriptome assembly were performed with Cufflinks v2.2.1 (Trapnell C. et al 2010) (RRintellectual disability:SCR_014597) using the relative UCSC mm10 GTF file. For each sample Cufflinks assembles the RNA-Seq reads into individual transcripts, inferring the splicing structure of the genes and classified them as known or novel. The output GTF files from each of the Cufflinks analysis and the GTF annotation file were sent to Cuffmerge v2.2.1 (Trapnell C. et al 2010) (RRintellectual disability:SCR_014597) to amalgamate them into a single unified transcript catalog.

### Isoform expression analysis from RNAseq data

Transcript expression in fragments per kilobase of exon per million reads mapped (FPKM) were estimated and normalized using Cuffdiff v2.2.1 (Trapnell C. et al 2010) (RRintellectual disability:SCR_014597) with default parameters from Cuffmerge GTF file result and alignment files. The R package CummeRbund v2.16 (Trapnell C et all 2012) (RRintellectual disability:SCR_014568) was used for data exploration and figure generation of some isoforms of interest.

### Differential gene expression analysis

Gene counts were calculated with featureCounts v1.4.6-p4 (Liao Y et al., 2014) (RRintellectual disability:SCR_012919) from a GTF containing known UCSC mm10 genes as well as the novel genes detected by Cufflinks (Cufflinks class code “u”) and alignment files. The R package DESeq2 v1.14.1 (Love MI et al., 2014) (RRintellectual disability:SCR_000154) was then used to normalize counts and detect the differentially expressed genes (FDR < 0.05). Batch effect between replicates was added in the design formula of DESeqDataSetFromMatrix function to model it in the regression step and subtract it in the differential expression test.

### Data access

The RNA-Seq data discussed in this publication have been deposited in NCBI’s Gene Expression Omnibus and are accessible through GEO Series accession number GSE110491 (https://www.ncbi.nlm.nih.gov/geo/query/acc.cgi?acc=GSE110491).

### Gene ontology

Gene ontology enrichment was performed using all of the expressed genes as background. We used DAVID (RRID:SCR_003033) with high stringency parameters, and ClueGo (Cytoscape) with a similar approach. DAVintellectual disability adjusted p-values were used for further evaluation.

### RNA-Seq statistics

We assumed that the samples were normally distributed. *P*-values for overlaps were calculated with binomial test using a custom-made R script. *P*-values were subsequently adjusted for multiple comparisons using Benjamini-Hochberg FDR procedure. Two-way permutation test of 1000 was adapted to validate the overlaps. We randomized the differentially expressed gene sets by randomly selecting same number of genes from RNA-seq expressed genes and subsequently calculating the overlap P-values. Moreover, we adapted a permutation test to evaluate the detected differentially expressed genes, randomizing 1000 times the RNA-seq data and recalculating the differentially expressed genes. Analysis for RNA-seq was performed using custom made R scripts implementing functions and adapting statistical designs comprised in the libraries used. The heatmap and Volcano plots for gene expression was performed from the gene overlap file using scripts written on R.

### Coexpression network analysis

RPKM data from [ref 51] was obtained from 11 neocortical areas in all developmental time points and genes corresponding to module 37 were selected. We performed pairwise Pearson correlation in R for all selected genes on log2 + 1 transformed RPKM and calculated degree centrality for each gene when correlation was at least 0.70. Degree centrality and network visualization was performed with Cytoscape. We colored genes according to SFARI score (release 12-05-2019).

### Behavior

All behavioral tests were done with age-matched male 8 to 14 weeks old littermates. They were performed according to the European Union and national recommendations for animal experimentation. The experimenter was blind to the genotypes during all analyses.

#### Open field test

The test was performed in a 40 x 40 cm square arena with an indirect illumination of 100 lux. Mouse movements were video-tracked using Smart 3.0 software (Panlab, Harvard apparatus) for one hour. Total distance traveled, time in center (exclusion of a 5 cm border arena), resting time and mean velocity were measured. The open-field arena was cleaned and wiped with H20 and 70% ethanol between each mouse. Data shown are means +/− s.e.m. and were analyzed using one way ANOVA or Kruskall-Wallis ANOVA when required. (WT : n=16; HET : n=15; KO : n=15).

#### Stereotypic behavior

During the first 10 minutes of the open-field test, the number of rearing and circling were measured manually. Both on-wall and off-wall rearing were counted. An on-wall rearing event was counted when both front-paws were apposed on the wall. An off-wall rearing event was counted when both front paws had left the floor. A complete 360-degree turn of nose angle with respect to the mouse body center was counted as one circling event. Data shown are means +/− s.e.m. and were analyzed using Kruskall-Wallis ANOVA. (WT : n=16; HET : n=15; KO : n=15).

#### Three-chamber test

The three-chamber apparatus consisted of a Plexiglas box (60 x 40 cm, each chamber being 20 x 40 cm) with removable floor and partitions dividing the box into three chambers with 5-cm large openings between chambers. Test mice were housed individually the day before the test. The task was carried out in five trials of 5 min each. After each trial, the mouse was returned to his home cage for 15 min. The three-chambers apparatus was cleaned and wiped with 70% ethanol between each trial.

In the first trial, a test mouse was placed in the center of the three-chamber unit, where two empty wire cages were located in the left and right chambers to habituate the test mouse. The mouse was allowed to freely explore each chamber. The mouse was video-tracked for 5 minutes with Smart 3.0 software. In the second trial, an 8-weeks old C57Bl/6J mouse (M1) was placed randomly in one of the two wire cages to avoid any place preference. The second wire cage remained empty (E). The test mouse was placed in the center, and allowed to freely explore the chamber for 5 min. In the following two trials, the same mouse M1 was again placed in one of the wire cages, and the test mouse was placed in the center and allowed to explore each chamber. The goal of these 2 trials was to familiarize the test mouse with M1. In the last 5-min session, a new 8-weeks old C57Bl/6J mouse (M2) was placed in the second wire cage. Thus, the test mouse had the choice between a now familiar mouse (M1) and a new, stranger mouse (M2). Time spent in each chamber and time spent within a 5-cm square proximal to each wire cage with the nose towards the cage (that we called investigation time) were measured. All data presented are means +/− s.e.m. and analyzed using two-way ANOVA with Bonferroni’s post hoc analysis. (WT : n=14; HET : n=15; KO : n=13).

#### New object recognition test

The arena used for the novel object recognition test was the same used for the open-field test. The arena was cleaned and wiped with 70% ethanol between each mouse. For habituation, the tested mouse was placed in the arena and allowed to explore for 10 min. Following habituation, two identical objects (50 ml orange corning tube) were placed in opposite corners of the arena, 10 cm from the sidewalls. The tested mouse was placed in the center of the arena, and allowed to explore the arena for 10 min. After 24 h, one object was replaced with another novel object, which was of similar size but differed in shape and color with the previous object (white and blue LEGO® bricks). The test mouse was placed in the center, and allowed to explore the arena and the two objects (a new and an “old” familiar object) for 10 min. The movement of the mice was video-tracked with Smart 3.0 software. Time in each proximal area (nose located in a 2 cm area around the object) was measured. All data shown are means +/− s.e.m. and analyzed using Student’s two-tailed, paired t-test or Wilcoxon Signed Rank Test when required. (WT : n=16; HET : n=15; KO : n=15).

### *In vitro NEUROD2* pathogenicity test

P19 cells culture and transfection were performed with modification from Farah et al., 2000 (*20*). P19 mouse carcinoma cells were cultured in MEMα with 7.5% bovine serum and 2.5% fetal bovine serum (HyClone) containing Glutamax and essential amino acids (Gibco), and maintained subconfluent prior to transfection. For lipofectamine 2000 (Invitrogen) transfections, cells were plated on glass coverslips in 24-well dishes at a concentration of 50,000 cells per well and transfected 24 hours later following manufacturer’s instructions. Three plasmids were co-transfected in each condition: pCAG-Cre, pCALNL-DsRed and pCALNL-*NEUROD2, NEUROD2* being either the WT form or any of the variants (see Fig. 1g-i). Cells were fixed 3 days post-transfection, immunostained for β3-tubulin (BioLegend BLE801201, 1:1000), counterstained with Hoechst solution (Molecular Probes) and mounted. For each condition we analyzed 3-4 independent experiments with 3 coverslips each.

**Figure 1:**
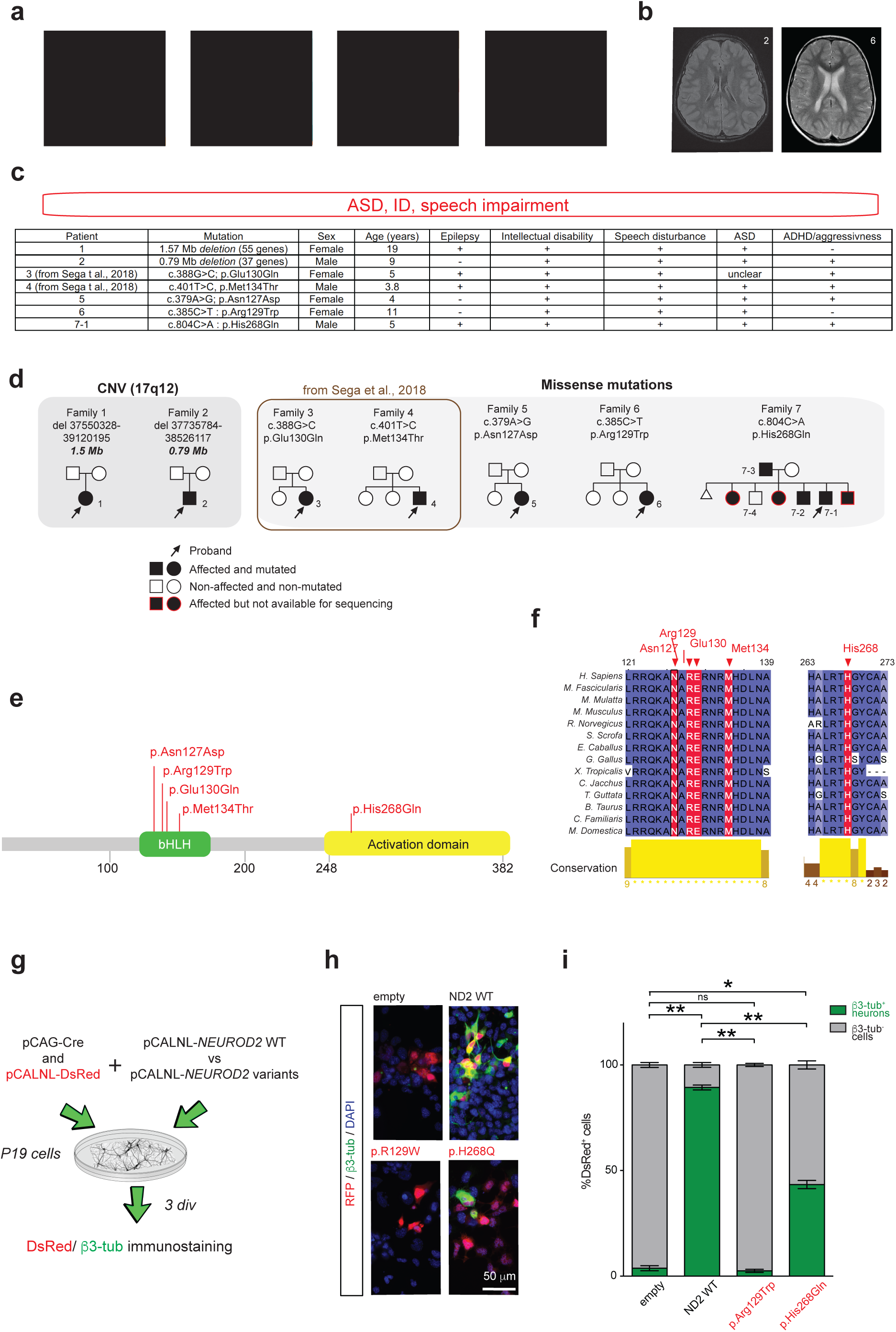
*NEUROD2* pathogenic mutations cause an ASD and intellectual disability syndrome. (**a-c**) The clinical phenotype of *NEUROD2* syndrome. (**a**) Affected individuals showed slightly dysmorphic facial features (shown are pictures from individuals in families 1, 6 and 7). (**b**) Axial T2-weighted MRI images of patients 2 and 6 showing no obvious abnormality in brain organization. Only patient 3 showed a thinned corpus callosum. (**c**) Main clinical characteristics of *NEUROD2* syndrome patients (see Supplementary Table 1 for more detailed information). (**d**) Pedigrees of 7 families with *NEUROD2* disruptions (Family 1 and 2 with copy number variations, Families 3 to 7 with 5 different pathogenic missense heterozygous mutations). Patients 7-1 and 7-2 in family 7 share the same mutation with their father (patient 7-3). Sanger sequencing showed that their healthy brother 6-4 does not carry the mutation in *NEUROD2*. (**e**) Localization of NEUROD2 mutations analyzed in this study, that are associated with intellectual disability/ASD. (**f**) Phylogenetic conservation of residues substituted in *NEUROD2* for the 5 patients with missense mutations. The positions of substituted residues are indicated in red. *In silico* multiple sequence alignment was performed using Clustal Omega and Jalview software. (**g-i**) Development of an *in vitro* test to measure the pathogenicity of *NEUROD2* variants. (**g**) P19 cells were transfected with pCAG-Cre, pCALNL-RFP and a plasmid containing different variants of Cre-dependent human *NEUROD2* (pCALNL-*NEUROD2*). (**h**) Representative images of RFP-transfected neurons immunostained for β3-tubulin (green) after 3 days *in vitro*. (**i**) The percentage of RFP^+^ cells expressing β3-tubulin in each condition was counted (n=3 independent experiments). Data are represented as means ± SEM. Statistical significance was evaluated by repeated measure one-way ANOVA followed by Sidak’s multiple comparisons post-hoc test (i) (\**P* < 0.05, \*\**P* < 0.01). Consent was obtained to publish the clinical images.

### Statistics

All statistical tests are described in the figure legends. All values represent the averages of independent experiments ± SEM. Statistical significance for comparisons with one variable was determined by Student’s t-test using two-tailed distribution for two normally distributed groups, and by Mann-Whitney test when distributions were not normal. Significance of multiple groups was determined by one-way, two-way, or two-way repeated measure ANOVA followed by either Bonferroni’s post hoc test as indicated in figure legends. In two-way ANOVA, the three assumptions for sampling data were tested: normality, homoscedasticity, independence. To test the assumption of residuals normality and homoscedasticity, Shapiro-Wilk test and Levene’s test were performed. If the assumption of normality was violated, a permutation test for a two-way ANOVA was used.

In particular, to statistically analyze the laminar distribution of the cells with Bins (Fig. 2), we fist checked if the variables were spatially autocorrelated in order to eliminate the effects of nearby spatial units. The purpose was to meet one of the conditions of two-way ANOVA: independence. We used spatial weights matrix and Moran’s I index to detect spatial autocorrelation. We chose a contiguity matrix as spatial weights matrix. If one Bin was connected with another Bin, the corresponding matrix was assigned the value 1, otherwise assigned the value 0. Then, the weights matrix was normalized by dividing each matrix line by its total. If the Moran’s test was not significant, the residuals in the traditional two-way ANOVA were not significantly spatially autocorrelated so a traditional two-way ANOVA could be performed. In the opposite situation, we filtered out the influence of spatial autocorrelation by applying the spatial lag model. The spatial autocorrelation parameter ρ in spatial lag model was estimated by using a maximum likehood method. In spatial lag model, the spatial autocorrelation in residuals was eliminated and therefore spatially adjusted two-way ANOVA can be performed.

**Figure 2:**
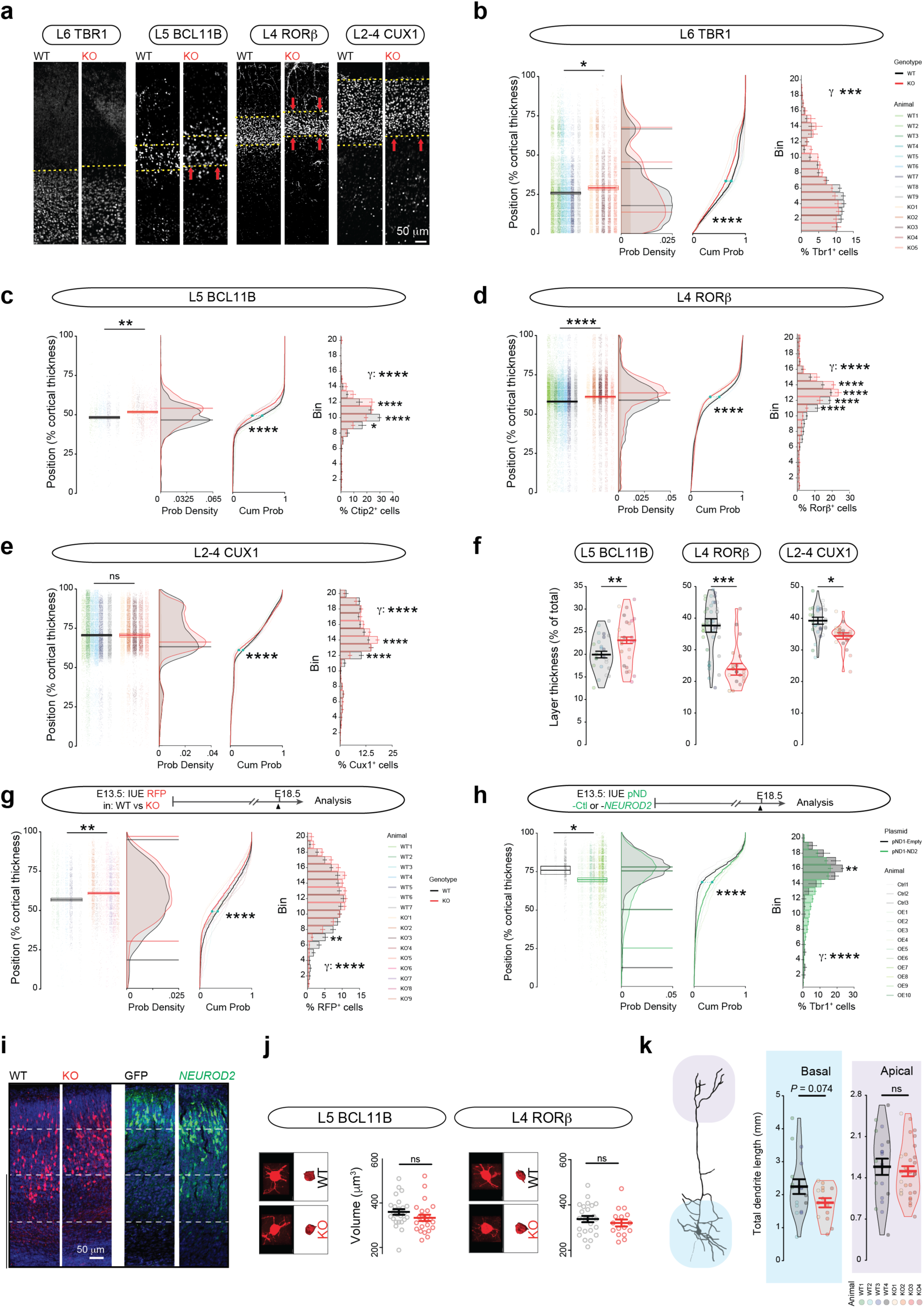
Altered laminar position and excess radial migration of CPNs in *Neurod2* null mice. (**a-f**) Altered laminar positioning of CPNs at P30. (**a**) Confocal photomicrographs of cortical columns at the somatosensory level. (**b-e**) Laminar positioning of CPN from different layers. Mean cell laminar position in percentage of the cortical thickness, probability density function (bars representing maximal probability density, for TBR1 the three peaks are represented), empirical cumulative distribution function and 20-bin based distribution of CPN types using the markers TBR1 for L6 (**b**), BCL11B for L5 (**c**), RORβ for L4 (**d**) and CUX1 for L2-4 combined (**e**). All layers were switched superficially, with the strongest significance for L5 and L4. (**f**) Layer thickness was increased for L5 and decreased for L4 and L2-4 (not measured for TBR1-expressing L6 because its upper border is not clearly identifiable). Data are shown as violin plot histograms where each circle is a coronal section from a color-coded mouse (statistics are calculated on the n of mice). (g-i) NEUROD2 limits radial migration of L5 CPNs. (**g**) L5 CPNs stopped their migration more superficially in *Neurod2* null mice that in WT littermates. Bars in the probability density function show start and end of cell density enrichment (max probability is equivalent between genotypes here). (**h**) Post-mitotic overexpression of *NEUROD2* in L5 CPNs of WT mice reduced migration of electroporated cells. Bars in the probability density function represent maximal probability densities in each quartile of the cortical thickness. (**i**) Representative confocal images of electroporated cells in *Neurod2* KO (red) or *NEUROD2*-overexpression (green) condition at E18.5. (**j,k**) Somatic volume and dendritic complexity were not significantly altered. Somas and dendrites were 3D-reconstructed with Imaris software. Data are represented as means ± SEM. Statistical significance was evaluated by Student t-test or Mann-Whitney test depending on normality of samples [Y cell position in (b)-(e) and (d)-(h), (j), (k)], by Kolmogorov-Smirnov test [cumulative probability graphs in (b)-(e) and (d)-(h)] or by Permutation test for a spatially adjusted two-way ANOVA followed by Bonferroni’s post-hoc test [bin graphs in (b)-(e) and (d)-(h)] (\**P* < 0.05, \*\**P* < 0.01, \*\*\**P* < 0.001, \*\*\*\**P* < 0.0001).

RNA-seq statistics are described above. Differences were considered significant when *P* < 0.05. All statistical analyses were performed with Prism 6 software (Graphpad).

## Results

### *NEUROD2* pathogenic mutations cause ASD and intellectual disability

Through a collaborative effort, we gained access to 7 families with pathogenic *NEUROD2* mutations associated with ASD, intellectual disability and speech delay. Among these 7 families, two had small *de novo* 17q12.3-q21 interstitial deletions (1.57 Mb and 0.79 Mb) including *NEUROD2 (*families 1 and 2, Fig. 1a-c, Supplementary Fig. 1) plus some other genes however unlikely to contribute to the phenotypes (Supplementary File 1 for the list of genes in deletions and their association with CNS function and disease). Two families (3 and 4) are followed up from previously published patients carrying *de novo* heterozygous missense *NEUROD2* mutations who have had early epileptic encephalopathic seizures from 5 to 16 month-old (*11*). Families 5 and 6 carry newly-identified *de novo* heterozygous missense *NEUROD2* mutations (c.379A>G; c.385C>T). Finally, family 7 is a non-consanguineous family with a *NEUROD2* heterozygous missense mutation i(c.804C>A) inherited from the affected father to his affected children but not to the only unaffected child (6/8 individuals affected; Fig. 1a-c). All affected patients had grossly normal brain MRIs (Fig. 1b), slightly dysmorphic facial features (Fig. 1a) and presented clinically with a spectrum of phenotypes including ASD, intellectual disability and speech disturbance (Fig. 1c). Comorbidities included epilepsy (3/7) and frequent hyperactivity/ADHD (5/7) (Fig. 1c).

Patients 3 to 7-3 with missense mutations were identified by whole-exome sequencing and confirmed by family-based Sanger sequencing. The 5 *NEUROD2* mutations from these patients were not found in 141,456 individuals assessed in GNOMAD. Of note, no other variants besides *NEUROD2* were identified in known disease-causing genes that could account for the clinical phenotypes in any of the 7 cases.

*De novo* mutations Glu130Gln, Met134Thr, Asn127Asp and Arg129trp from families 3-6 are in the conserved DNA-binding domain, suggesting pathogenicity (Fig. 1e,f). In contrast, the His268Gln mutation in the familial case lies in the activation domain, which is also conserved (Fig.1f).

The pathogenicity of Glu130Gln and Met134Thr mutations from families 3 and 4 has already been reported (*11*). To test the pathogenicity of novel Arg129trp and His268Gln mutations found in patients 6 and 7-1 to 7-3, we took advantage of the fact that a WT human *NEUROD2* plasmid transfected in P19 embryonic carcinoma cells induces the rapid differentiation of these cells into β3-tubulin expressing neurons [63] (Fig. 1g). While human WT *NEUROD2* was able to induce neuronal differentiation in ∼ 90% of transfected cells, the Arg129trp and His268Gln mutations were not and less able to direct neuronal differentiation, respectively (∼2.5 and 35% of transfected cells, respectively; Fig. 1h,i). These data indicate that the Arg129trp and His268Gln *NEUROD2* mutations are pathogenic and probably loss-of-function.

### NEUROD2 is restricted to projection neurons in the embryonic and postnatal mouse neocortex

To identify cell types that express *Neurod2* in the neocortex, we first performed quantitative real time PCR from whole cortex of embryonic and postnatal stages. *Neurod2* mRNA expression increased from E14.5 to reach a peak at E18.5, and remained at lower albeit constant levels postnatally (Supplementary Fig. 2a). In situ hybridizations from ALLEN BRAIN ATLAS (http://www.brain-map.org/) revealed *Neurod2* mRNA expression in the cortical plate and hippocampus starting from E13.5 onwards (Supplementary Fig. 2b). In the postnatal cortex, *Neurod2* mRNA was maintained after birth, and observed in L2 to L6 at all ages examined.

At the protein level, NEUROD2 was also detected embryonically, in the intermediate zone where migration occurs and in the cortical plate but not in the germinal ventricular /subventricular zone (V-SVZ, Supplementary Fig. 2c), confirming results of a previous study (*21*). NEUROD2 protein was maintained postnatally in cortical L2 to L6 (Supplementary Fig. 2c). It co-localized with the L5 pyramidal tract CPN marker BCL11B (Supplementary Fig. 2d,e) and with other CPN markers that represent other layers (not shown) at all ages examined (P3, P30, P90 and 1 year, not shown). In contrast, NEUROD2 was never detected in Gad67-GFP^+^ inhibitory neurons (Supplementary Fig. 2e), nor in non-neuronal cortical cells (not shown). In sum, *Neurod2* mRNA and NEUROD2 protein are expressed in migratory and post-migratory CPNs of L2 to L6.

### *Neurod2* deletion alters relative size and positioning of cortical layers

To gain insight into the pathophysiological mechanisms underlying ASD and intellectual disability in humans, we performed a systematic and multi-level analysis of cortical development in *Neurod2* KO mice.

First, we analyzed the gross anatomy of the *Neurod2* KO cortex at P30, when CPNs should have stopped migrating and be fully integrated into the cortical circuit. The corpus callosum of *Neurod2*-null mice showed a slightly reduced thickness (Supplementary Fig. 2f), but global cortical anatomy appeared otherwise normal, as indicated by the normal aspect and thickness of the cortical wall at sensory-motor levels (Supplementary Fig. 2g).

The specialization of CPN subtypes is largely determined by their axonal outputs (*22–24*). Thus, we asked if axonal targeting specificities of CPN subtypes were altered in absence of *Neurod2*. We injected the retrograde tracer cholera toxin beta (CTB) at each of the three main axonal target regions of motor and somatosensory cortices: the ventrolateral thalamus targeted by L6 cortico-thalamic PNs, the striatum preferentially targeted by a subset of L5 CPNs, and L2-5 of M1 targeted by contralateral L2-5 callosal CPNs. Thalamic, striatal and M1 injections retrogradely labeled CPN somata in L6, L5 and contralateral L2-5, respectively (Supplementary Fig. 2h,i), like in WT littermates. Labeling major forebrain axonal tracts using L1 immunostaining confirmed normal organization of major axonal tracts (Supplementary Fig. 2g). Thus, axonal targeting specificities of CPN subtypes were normal in *Neurod2* KO mice.

We then quantified the amount and laminar distribution of CPN subtypes in S1 and M1 using immunostainings for major regulatory transcription factors of selected cortical layers. We used TBR1 for L6, BCL11B for L5 pyramidal tract, RORβ for L4 and CUX1 for L2-4 CPNs. Each CPN type was found in normal quantities in a cortical column spanning the width of the cortex (Supplementary Fig. 2j), indicating normal production of CPNs and thus unaltered proliferation and apoptosis of CPN precursors during development. This was to be expected given that NEUROD2 is not expressed in VZ progenitors. Moreover, the inside-out patterning was normal since TBR1, BCL11B, RORβ and CUX1 were successively expressed from deep to superficial layers (Fig. 2a). However, interestingly the upper borders of TBR1 and BCL11B-expressing deep layers extended superficially towards the pial surface in *Neurod2* KO mice (Fig. 2b,c). This superficial extension of L5-6 was correlated with a significant enlargement of L5 towards the brain surface (Fig. 2a,c,f) and a significant thinning of RORβ and CUX1-expressing superficial L2-4 that were also switched superficially (Fig. 2d-f). L2-4 were squeezed between L1 and the enlarged L5, hence leading to an overall unaltered cortical thickness (Supplementary Fig. 2g).

### *Neurod2* deletion induces excessive radial migration of CPNs during embryogenesis

The increased *vs* decreased surfaces occupied by L5-6 *vs* L2-4 in *Neurod2*-null mice can result from altered radial migration or from post-migratory maturational defects such as changes in cell size for some CPN types. To discriminate between these two hypotheses, we first analyzed the radial migration of L5 CPNs using *in utero* electroporation. A red fluorescent plasmid was electroporated at E13.5 when cortical progenitors generate L5 CPNs, and embryos were sacrificed around the end of radial migration, at E18.5. Analyses including mean cell laminar position, probability density, cumulative distribution and bin-based laminar distribution all demonstrated that in KO mice electroporated L5 CPNs had reached more superficial levels at E18.5, demonstrating over-migration (Fig. 2g,i). Conversely, overexpressing *Neurod2* in post-mitotic CPNs of WT mice using the *NeuroD1* promoter increased the fraction of electroporated cells in deep layers at E18.5 compared to a control group (Fig. 2h,i). Taken together, these data indicate that *Neurod2* cell autonomously controls the precise laminar positioning of CPNs by limiting the distance of their radial migration during corticogenesis. Hence, the altered layer positioning in *Neurod2* KO mice is, at least in part, due to radial migration defects in CPNs during embryogenesis.

Because post-migratory alterations in cell or neuropil volumes could also contribute to the respective enlargement and thinning of L5 and L4, we assessed somatic volumes and dendritic morphologies. Soma volume at P30 measured from 3-D volumetric reconstructions of RFP-electroporated CPNs did not vary in L5 and L4 (Fig. 2j). We measured the complexity of integral L5 CPN dendritic arbors by using the Thy1-GFP mouse, line M where GFP levels reach levels in fine dendritic processes far higher than the ones obtained with electroporations (*25*). At P30, the overall size of basal and apical dendrites of L5 CPNs was not significantly altered in *Neurod2* null mice (Fig. 2k). Overall, our data strongly suggest that the layer positioning phenotype in *Neurod2* KO mice is a consequence of radial migration defects but not of post-migratory variations in neuronal sizes.

### *Neurod2* deletion alters dendritic spine density and turnover in CPNs

*Neurod2* expression in post-migratory CPNs is consistent with the hypothesis that this transcription factor can also play a regulatory role on synaptogenesis (*26*). The Thy1-GFP mouse line M allowed us to gained experimental access to the fine morphology of dendritic spines (*25*), the post-synaptic elements of excitatory synapses. We measured spine density on basal and apical compartments at two ages, i.e. P30 and P120 (corresponding to juvenile and adult ages, respectively). In basal dendrites, spine density was slightly reduced at P30 but had reached WT levels at P120 (Fig. 3a,b and Supplementary Fig. 3a,b). Overall the temporal evolution of dendritic spine density in basal compartments was similar to the WT situation, with a strong reduction between P30 and P120 in both genotypes that reflected spine pruning (Fig. 3b). The phenotype in the apical tuft was more pronounced. There, spine density was more strongly reduced at P30 and it was not significantly reduced with aging, contrasting with the WT situation (Fig. 3c). At P120, spine density in KO actually tended to increase compared to WT (*P*= 0.07).

**Figure 3:**
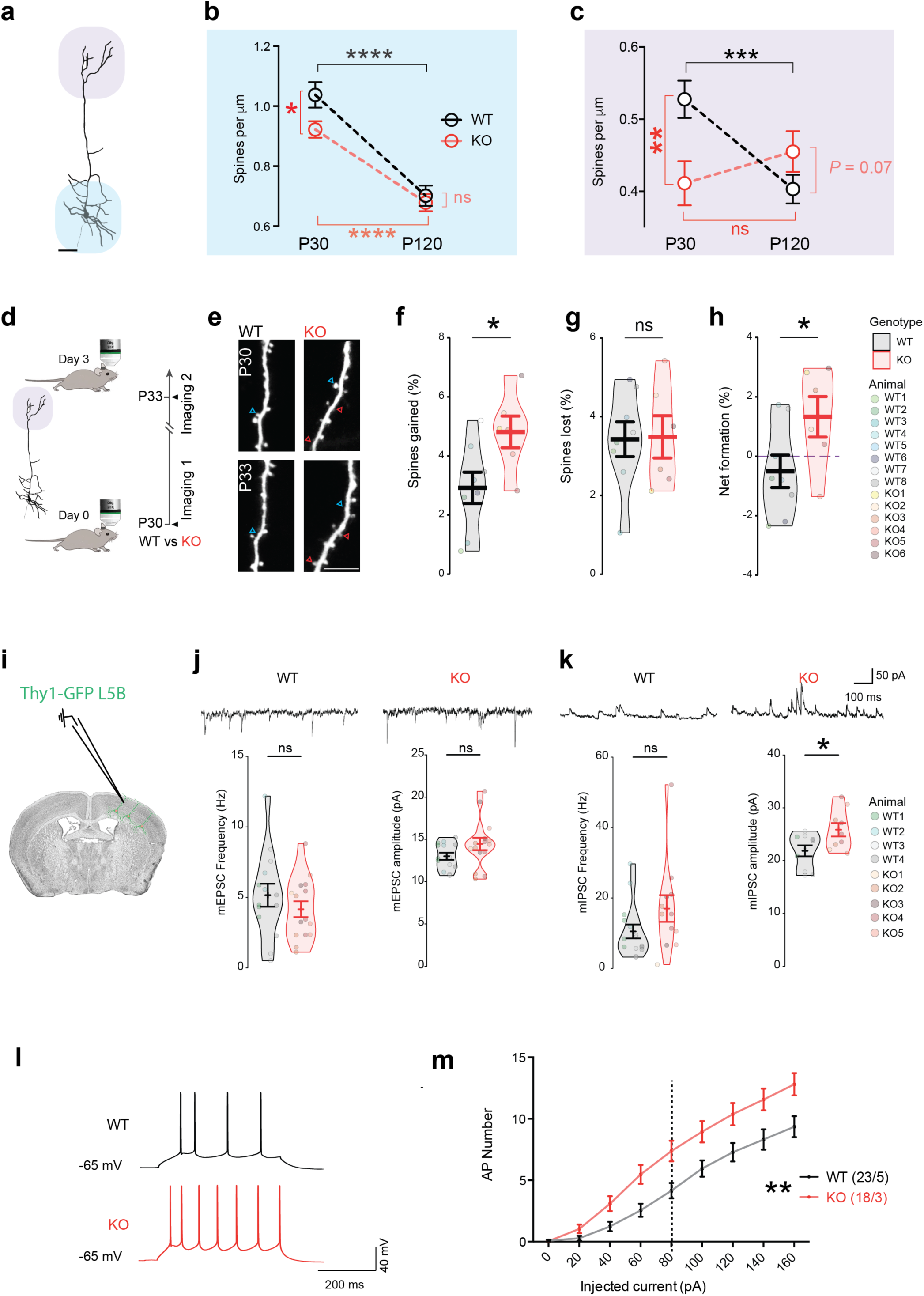
Synaptic and physiological defects in CPNs of *Neurod2* KO mice. (**a-c**) Age-dependent spine density defects in basal and apical dendrites. (**a**) Imaris reconstruction of an L5 CPN, basal and apical compartments are underscored by blue and purple halo, respectively. (b,c) Spine density at P30 and P120 in basal (**b**) and apical (**c**) compartments. (**d-h**) Increased spine turnover in apical tuft. (**d**) Experimental paradigm. (**e**) Representative 2-photon images of same dendrites at 3 days interval. Red and blue arrowheads depict gained and lost spines, respectively. (**f**) Spine gain, (**g**) spine loss and (**h**) net spine formation over the 3-day interval period shown in violin plot histograms (each circle is an animal). (**i-k**) Miniature post-synaptic currents. (**i**) Patch clamp recording of Thy1-GFP CPNs in L5B. (**j**) Representative traces, frequency and amplitude of mEPSCs (each circle is a recorded cell with a mouse color code in the violin plots). (**k**) Example traces, frequency and amplitude of mIPSCs. Amplitude of mIPSC was significantly increased. (**l,m**) Increased intrinsic excitability. (**l**) Representative firing responses to +80 pA current steps in a WT (black) and a KO (red) cell. (**m**) *Neurod2* KO neurons reached action potential (AP) firing threshold earlier than WT neurons and exhibited a steeper input-output relationship, as assessed by the number of APs elicited by increasing current injections (from +20 to +160 pA, 20-pA increments) during current-clamp recordings. Data are represented as means ± SEM. Statistical analyses were performed using two-tailed t-tests or Mann-Whitney test depending on the normality of samples [(f) to (k)], by two-way ANOVA followed by Bonferroni’s post hoc test [(b), (c)] and by two-way repeated measure ANOVA followed by Bonferroni’s post hoc test [(m)]. \**P* < 0.05, \*\**P* < 0.01.

These data suggest that spine remodeling and turnover in L5 CPN apical tufts might be abnormal from P30 onward in *Neurod2* mutant mice. We used 2-photon live imaging to measure this parameter in M1 (Fig. 3d). We counted the number and proportion of gained and lost spines on single identified dendritic segments at 3 days interval between P30 and P33. Gained and lost spines compensated for each other in WT mice (Fig. 3e-g). In contrast, gained spines outnumbered lost spines in *Neurod2* KO mice, resulting in a net spine gain over the 3-day observation period (Fig. 3e-h). In sum, spine density was still increasing at P30 in *Neurod2* KO mice, in contrast to WT mice where it had reached a plateau. Hence, the development of excitatory synapses is altered in L5 CPN apical tufts of *Neurod2* KO mice.

### Increased amplitude of inhibitory inputs and excitability in L5 CPNs of *Neurod2* KO mice

The lamination phenotype as well as dendritic spine alterations observed in L5 CPNs at P30 prompted us to study the electrophysiological properties of these cells. The frequency and amplitude of somatic miniature excitatory post-synaptic currents (mEPSCs) were unaltered (Fig. 3j). When measuring miniature inhibitory post-synaptic current (mIPSCs) we found no change in frequency but an increased amplitude in *Neurod2* KO mice (Fig. 3k). Because L2/3 CPNs receive reduced mEPSC and mIPSC frequencies but normal amplitudes in absence of *Neurod2* (*16*), we conclude that synaptic inputs are distinctly altered depending on the layer in *Neurod2* KO mice.

Next, we measured intrinsic physiological properties. Resting membrane potential as well as action potential threshold and amplitude were normal in *Neurod2* KO mice (Supplementary Fig. 3c-e). However, input membrane resistance was slightly increased (Supplementary Fig. 3f) and capacitance showed a trend towards reduction (Supplementary Fig. 3g). More strikingly, compared with WT, *Neurod2* KO L5 CPNs fired significantly more action potentials in response to depolarizing current injections (Fig. 3l,m), demonstrating increased intrinsic excitability. This phenotype was neither due to variations in action potential after-hyperpolarization (Supplementary Fig. 3h) - which contrasts with L2/3 CPNs in *Neurod2* deficient mutant (*16*) - nor to alterations in resting membrane potential or action potential threshold and amplitude (Supplementary Fig. 3c-e). Finally, we measured hyperpolarization-activated cation (HCN) Ih currents since they influence intrinsic neuronal excitability (*27*) and are critical integrators of synaptic integration selectively in L5 CPNs (*28*). Compared with WT mice, L5 CPNs in *Neurod2* KO mice exhibited a significantly increased Ih current density (Supplementary Fig. 3i,j), suggesting that *Neurod2* deficiency alters the expression or function of HCN channels. Since increased Ih current usually down-tunes intrinsic neuronal excitability (*27*), this phenotype might represent a compensatory mechanism that counter-balances *Neurod2*-induced neuronal hyper excitability.

### *Neurod2* deletion alters the expression of neuronal voltage-gated ion channels

To identify genes that are regulated by *Neurod2* in the neocortex, we performed RNA-seq on motor plus somatosensory cortex isolated from *Neurod2* WT versus KO mice at P30. Comparative analysis of RNA-seq identified 264 differentially expressed genes, among which 185 were downregulated and 79 were upregulated in KO mice (Fig. 4a,b; Supplementary File 1). To determine whether some of the differentially expressed genes are expressed in neuronal subtypes and might have functional roles in their development, we integrated information from the literature and several public data sources (DeCoN, Allen Brain Atlas, Single Cell Analysis of Mouse Cortex, GenePaint, Eurexpress). This search yielded 171 genes generic to all layers and 83 genes which had layer specificity (Fig. 4c,d). The proportion of layer-specific factors among differentially expressed genes was similar in L2/3, L5 and L6 (Fig. 4c,d), consistent with the idea that *Neurod2* impacts CPNs from all layers. qRT-PCR for 10 differentially expressed genes representative of different expression profiles (*Kcnh1* and *Cdh8* for L2/3, *Kcnq5* and *Htr2a* for L5, *Scn4b*, *Kcnk4*, *Scn8a*, *Scn1a*, *Cacna1c* and *Grin2b* for L2-6) validated the RNA-seq results by showing 100% concordance (Fig. 4e).

**Figure 4:**
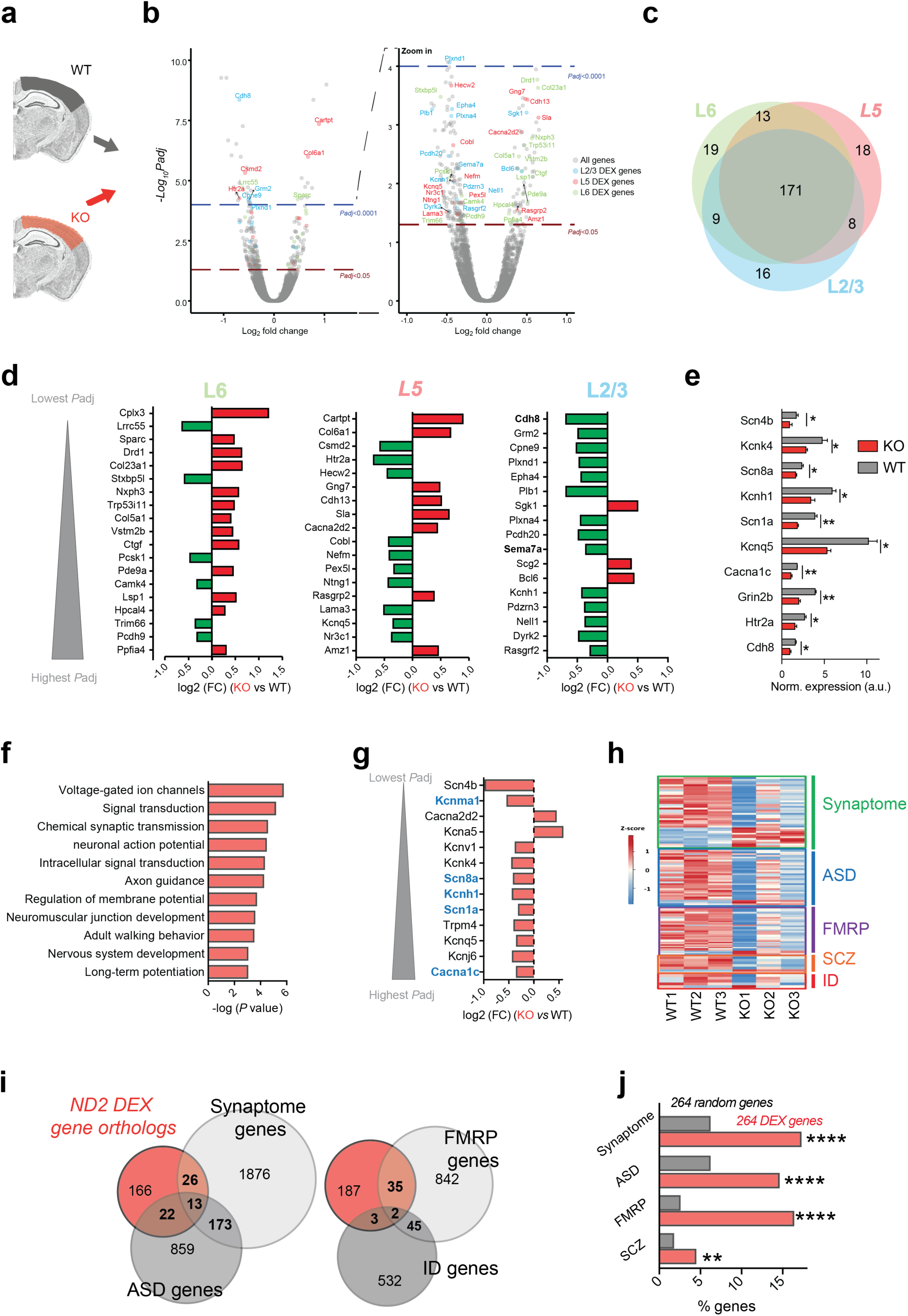
*Neurod2* KO mice show altered gene expression of cortical layer markers, voltage gated ions and ASD-related genes. (**a**) Schematic of the RNA-seq experiment, performed at P30. (**b**) Volcano plots depicting differential gene expression (N=3 experiments). (**c**) Venn diagram of differentially expressed genes specific to L2/3, L5 and L6 (171 are generic to all layers, some are common between 2 layers and others specific to a single layer). (**d**) Fold change expression (FC; log2 scale) of differentially expressed genes belonging to specific layers, ranked according to adjusted *P* value (lowest *P*adj value on top). (**e**) Validation of RNA-seq data by qPCR for 10 ion channel/ neuropsychiatric-related genes. N=3 experiments per genotype. (**f**) Gene set enrichment analysis using the DAVID knowledgebase. (**g**) Fold change expression (FC; log2 scale) of 13 differentially expressed genes belonging to the voltage-dependent ion channel family, ranked according to *P*adj (lowest on top). Genes associated with neuropsychiatric recurrent syndromes are depicted in blue. (**h-j**) Association between the human orthologs of differentially expressed genes and neuropsychiatric diseases. (**h**) Heatmap for synaptic and disease gene sets among the orthologs of differentially expressed genes (red, higher expression; blue, lower expression). (**i**) Venn diagrams identifying overlaps between differentially expressed genes and synaptic/ neurodevelopmental disease gene sets. Number of genes for each gene set is indicated. (**j**) Graphical representation of % [differentially expressed genes] (orange) and % [same number of randomly selected genes] (grey) belonging to gene sets of interest. Statistical significance was evaluated by two-tailed Student t test (e) or binomial test (j) (*P < 0.05; **P < 0.01; ****P < 0.0001).

Gene ontology (GO) analysis with DAVID (Fig. 4f and Supplementary Fig.4a) and ClueGO (*29*) (Supplementary Fig. 4b) revealed that differentially expressed genes were most significantly enriched for voltage-gated ion channel activity, followed by cell projection morphogenesis and chemical synaptic transmission. We found 13 differentially expressed voltage-gated ion channels including the sodium channels *Scn1a*, *Scn4b* and *Scn8a*, the potassium channels *Kcnh1*, *Kcnq5*, *Kcnj6*, *Kcna5*, *Kcnv1*, *Kcnk4* and *Kcnma1*, and the calcium channels *Cacna1c* and *Cacna2d2* (Fig. 4g), many of which are associated with neuropsychiatric syndromes (*Scn1a* in Dravet syndrome, *Cacna1c* in Timothy syndrome, *Kcnma1* in developmental delay and epilepsy, *Scn8a* in early epileptic encephalopathy and cognitive impairment, *Kcnh1* in Temple-Baraitser syndrome; blue in Fig. 4g). Among those 13 differentially expressed ion channels, 11 were downregulated and 2 were upregulated (*Cacna2d2* and *Kcna5*). In summary, we find that a genetic network controlling neuronal excitability and post-synaptic development is regulated by *Neurod2* in the mouse P30 M1/S1 neocortex.

### *Neurod2* deletion modifies ASD-associated gene expression

Because 227 of the 264 mouse differentially expressed genes had a non-ambiguous human ortholog, we reasoned that examining their disease association could provide a valuable indication to *NEUROD2* function. Extensive PubMed searches for all human orthologs of differentially expressed genes established that 190 of the 227 genes (83.7%) are putative causal loci for brain and/or nervous system disorders. Interestingly, the great majority of these genes (146/190; 76.8%) have been associated with ASD, with the second most represented disease (5.8%) being schizophrenia (Supplementary File 1).

In a similar approach, we analyzed the overlap between the orthologs of differentially expressed genes and ASD-associated genes (*30*) from SFARI database (859 genes), FMRP-associated genes from Darnell and colleagues (*31*) (842 genes), intellectual disability-associated genes (*32*), and schizophrenia-associated genes ((*33*) and OMIM, 196 genes) or synaptome-associated genes (*34*) (SynaptomeDB, 1876 genes), and generated a heatmap of expression levels for overlapping genes (Fig. 4h). Venn diagrams showed that among the 264 differentially expressed genes, 39 (17.1%) were associated with the synaptome, 35 (15.4%) with ASD, 37 (16.3%) with FMRP and 4.4% with schizophrenia (Fig. 4i and Supplementary File 1). These proportions are much higher than what a random sampling would produce (Fig. 4j and Supplementary File 1 for binomial test with 1000 resamplings; hypergeometric test gave comparable significance). Differentially expressed ASD genes included the Dravet syndrome gene *SCN1A* and the Timothy syndrome gene *CACNA1C*, but also *GRIN2B*, *HTR2A*, *PCDH9* and many other genes (Supplementary Fig. 5). Among the synaptome-related differentially expressed genes, 4 were pre-synaptic and 36 were post-synaptic (*34*) (Supplementary Fig. 5), suggesting that *Neurod2* primarily regulates the post-synaptic element of synapses.

### *NEUROD2* is a nexus in a human brain developmental gene network whose defects are associated with ASD

We studied the localization of *NEUROD2* within the 73 gene co-expression modules obtained by weighted gene correlation network analysis (WGCNA) from human neocortical samples spanning almost the entire lifespan from psychENCODE (*35*). We found NEUROD2 in module 37, which is composed of 145 genes enriched in neuronal cell types and fetal enhancers. This module showed also exceptional enrichment in association signals from multiple neuropsychiatric traits and disorders, including schizophrenia, neuroticism and IQ (*35*), as well as in genes related to ASD from the analysis of de novo mutations, associated to developmental delay and/or described in SFARI. NEUROD2 presented very high degree centrality within module 37 (Supplementary Fig. 6a), and correlated expression with genes listed in SFARI with scores from 1 to 3, including *TCF4, TSHZ3*, *TBR1*, *NR4A2* and *SOX5* (Supplementary Fig. 6b).

Altogether, our molecular analyses in mice and humans point to *NEUROD2* as a node in a brain developmental gene network whose defects are associated with neurodevelopmental disorders including ASD.

### *Neurod2* loss and haploinsufficiency result in autism-like behaviors

We then investigated whether *Neurod2* loss or haploinsufficiency (that genetically mimic the patient condition) could result in ASD-like phenotypes by measuring the two core features, that is, impairment of social interactions and stereotyped repetitive behaviors, which serve to diagnose ASD (DSM-5).

The first criterion, that is, impaired social interactions, was evaluated using the three-chamber test. In the sociability task, unlike wild-type mice *Neurod2* KO mice did not interact more frequently with a conspecific than with an empty box (Fig. 6a-c and Supplementary Fig. 7a). *Neurod2* HET mice behaved similar to WT in this task, however (Fig. 6c). In the social recognition test, WT mice showed strong preference for interacting with the novel mouse, as expected (Fig. 6d-f and Supplementary Fig. 7b). In contrast, both *Neurod2* KO and HET mice did not interact more with the novel mouse (Fig. 6f, Supplementary Fig. 7b). Thus, social interactions were altered solely in *Neurod2* KO mice while social recognition/memory was impaired in both *Neurod2* KO and HET mice. In contrast to the social tasks, interest and memory for objects were unaltered in both genotypes, as both *Neurod2* HET and KO mice spent more time investigating a novel object in the novel object recognition task, similar to WT littermates (Fig. 6g-i). This confirms the specificity of the social behavior alteration.

The second ASD criterion, namely repetitive patterns of behavior, was assessed by analyzing ASD-related stereotypic mouse behaviors such as grooming, circling and rearing (*36*). Because mice of all 3 genotypes displayed almost no grooming behaviors owing to the mixed C57BL/6N;129Sv genetic background, we focused on rearing and circling. As previously described (*37*), *Neurod2* HET mice displayed increased circling (Fig. 6j). Instead, *Neurod2* null mice showed increased rearing, both at and outside the walls of the cage (Fig. 6j).

Finally, we assessed mouse behaviors related to ASD comorbidities, such as epilepsy, change in locomotor activity (ADHD-related phenotypes) and conflict anxiety. A third of *Neurod2* KO mice displayed spontaneous epileptic seizures observed at the behavioral level (Fig. 6k and Supplementary Video 1 for 2 examples), which were lethal for 3 out of 5 KO mice. One HET mouse had an epileptic crisis, while in contrast no WT mice showed behavioral seizure activity. The proportion of epileptic seizures among *Neurod2* KO and HET mice is likely an underestimate since mice were not monitored continuously. *Neurod2* KO mice were also hyperactive in the open field (Fig. 6l, Supplementary Fig. 7c). Indeed, they travelled significantly more than WT and HET littermates, while HET mice displayed only subtle hyper-locomotion compared to WTs (Fig. 6l). *Neurod2* KO mice displayed higher displacement velocities and shorter resting periods (Fig. 6l), confirming locomotor hyperactivity. Finally, we measured conflict anxiety, which requires functional integrity of the cerebral cortex (*38*) and is often altered in ASD (*39*). The open field arena presents a conflict between innate drives to explore a novel environment and safety. Under brightly lit conditions, the center of the open field is aversive and potentially risk-laden, whereas exploration of the periphery provides a safer choice. *Neurod2* HET and KO mice explored the center of the arena more often than their WT littermates (Supplementary Fig. 7d).

In summary, *Neurod2* HET and KO mice exhibited strong behavioral defects reminiscent of the symptoms (altered social interest and memory, stereotypies) and comorbidities (epilepsy, hyperactivity, reduced conflict anxiety) of ASD. *Neurod2* HET mice had milder phenotypes than *Neurod2* KO mice, indicating that *Neurod2* is a haploinsufficient gene.

### Forebrain excitatory neuron-specific *Neurod2* deletion recapitulates ASD-related phenotypes in mice

*Neurod2* expression is not restricted to the forebrain: it has been observed in paraventricular hypothalamic nuclei, cerebellum and spinal cord as well as in extra-CNS areas such as thyroid and pancreas (Fig. 7a). Importantly, paraventricular hypothalamic nuclei and thyroid have been associated with ASD. Hence, the phenotypes observed in full KO mice and *NEUROD2*-mutated patients might result from alterations in areas other than the forebrain. Furthermore, although we did not detect *Neurod2* mRNA and NEUROD2 protein in inhibitory neurons, it remains possible that *Neurod2* is expressed in these cell types at stages of their development that were not analyzed.

To directly evaluate the contribution of excitatory neurons of the forebrain to the ASD phenotypes, we generated conditional *Neurod2*^flox/flox^ mice in which the whole coding exon was flanked by loxP sites (Supplementary Fig. 8a), and crossed them with *Emx1*-Cre mice (*40*). Strikingly, forebrain excitatory neuron-specific *Neurod2* deletion recapitulated the major ASD-related behavioral defects, as both social interaction and recognition were strongly altered in *Emx1*-Cre; *Neurod2*^flox/flox^ mice compared to Cre-negative control littermates (Fig. 7b-e). In fact, quite surprisingly forebrain-specific *Neurod2* KO mice showed an even stronger phenotype in the 3-chamber test than full KO mice. In full KO mice the reduced mouse and stranger box investigations were compensated by increased investigations of empty and familiar box in the interaction and recognition test, respectively, so that the total overall interaction time was unchanged in both tests. In contrast, there was no compensation in forebrain-specific KO mice and the total interaction time (both boxes) was reduced (Fig. 7b,d). Forebrain excitatory neuron-specific *Neurod2* KO mice were also behaving normally in the novel object recognition task (Supplementary Fig. 8b) and hyperactive in the open field (Supplementary Fig. 8c), thus recapitulating the *Neurod2* full KO phenotype. We also observed spontaneous seizures in *Emx1*-Cre; *Neurod2*^flox/flox^ mice but not in control littermates, similar to the full KO situation (Supplementary Video 1).

The only difference between *Emx1*-Cre; *Neurod2*^flox/flox^ and full KO mice was in the time in open field center. While full KO spent more time, *Emx1*-Cre; *Neurod2*^flox/flox^ mice spent less time than controls in the center (Supplementary Fig. 8d). This indicates that other structures, likely the amygdala, that maintain their *Neurod2* expression in Emx1-Cre; *Neurod2*^flox/flox^ mice, are involved in the conflict anxiety phenotype We then aimed to test whether the over-migration observed in full *Neurod2* KO mice is cell autonomous by repeating the *in utero* electroporation strategy with Cre recombinase versus an empty plasmid in *Neurod2* cKO mice. L5 CPNs from cKO mice transfected with Cre recombinase indeed migrated excessively towards the pial surface (Fig. 7f), unambiguously demonstrating that the cell autonomous nature of the migration phenotype. This suggested that *Neurod2* deletion only in forebrain CPNs would be sufficient to induce the layer positioning phenotypes already described in *Neurod2* full KO mice. This was indeed the case, as forebrain excitatory neuron-specific KO mice showed reduced thickness and superficial switch of L4 at P30 (Fig. 7g,h and Supplementary Fig. 8e), reminiscent of the full KO phenotype.

## Discussion

In the present study, we identified the *NEUROD2* gene as responsible for a new neurodevelopmental syndrome. The characteristic clinical features include autistic traits, intellectual disability and speech disturbance. Mutations occurred de novo for 6/7 patients while it was inherited from an affected parent in a familial case, overall supporting the causative link between *NEUROD2* mutation and the syndrome. While a previous study has suggested that NEUROD2 is involved in early epileptic encephalopathy (*11*), our data point to a core NEUROD2-associated phenotype centered on ASD, intellectual disability and speech disturbance.

Non-fully penetrant *NEUROD2*-associated phenotypes include ADHD symptoms (5/7 patients) and epilepsy (3/7 patients). Importantly, our easy-to-perform and efficient *in vitro* NEUROD2-mediated neuronal differentiation assay could demonstrate the pathogenicity of *NEUROD2* variants (Fig. 1g) found in this study. Of interest for clinicians, testing the pathogenicity of novel *NEUROD2* variants will now be easy and rapid by using this *in vitro* assay. The causative link between *NEUROD2* deletion and the newly reported syndrome is strengthened by the mouse studies. Indeed, *Neurod2* mutation in the mouse recapitulates most features of the human phenotype: ASD-relevant social abnormalities and propensity to epilepsy and hyperactivity. Also, the expression profile of NEUROD2 reinforces the concept, emerging from single-cell sequencing data of the human brain (*41*), that mid-gestation and adult glutamatergic projection neurons are the most highly enriched cells in ASD genes. Notably, the DECIPHER human database reports a small, 135 kb-long duplication encompassing *NEUROD2* that is associated with intellectual disability and ADHD. This suggests that not only reduced but also increased expression levels of functional *NEUROD2* generate comparable symptoms, similar to what has been reported for the high confidence ASD genes *MECP2* (*42*) and *SHANK3* (*43*).

Our unbiased coexpression network analysis using psychENCODE (*35*) positioned *NEUROD2* in module 37, which shows an exceptional enrichment in association signals from multiple neuropsychiatric traits and disorders, including intellectual disability and ASD (*44*). *NEUROD2* presented a high degree centrality within this module (Fig 5a), and correlated expression with high confidence ASD genes listed in SFARI, including *TCF4, TSHZ3*, *TBR1*, *NR4A2* and *SOX5* (Fig 5b). These data provide support for a hub position of *NEUROD2* in a cortical transcriptional regulatory network associated with ASD.

**Figure 5:**
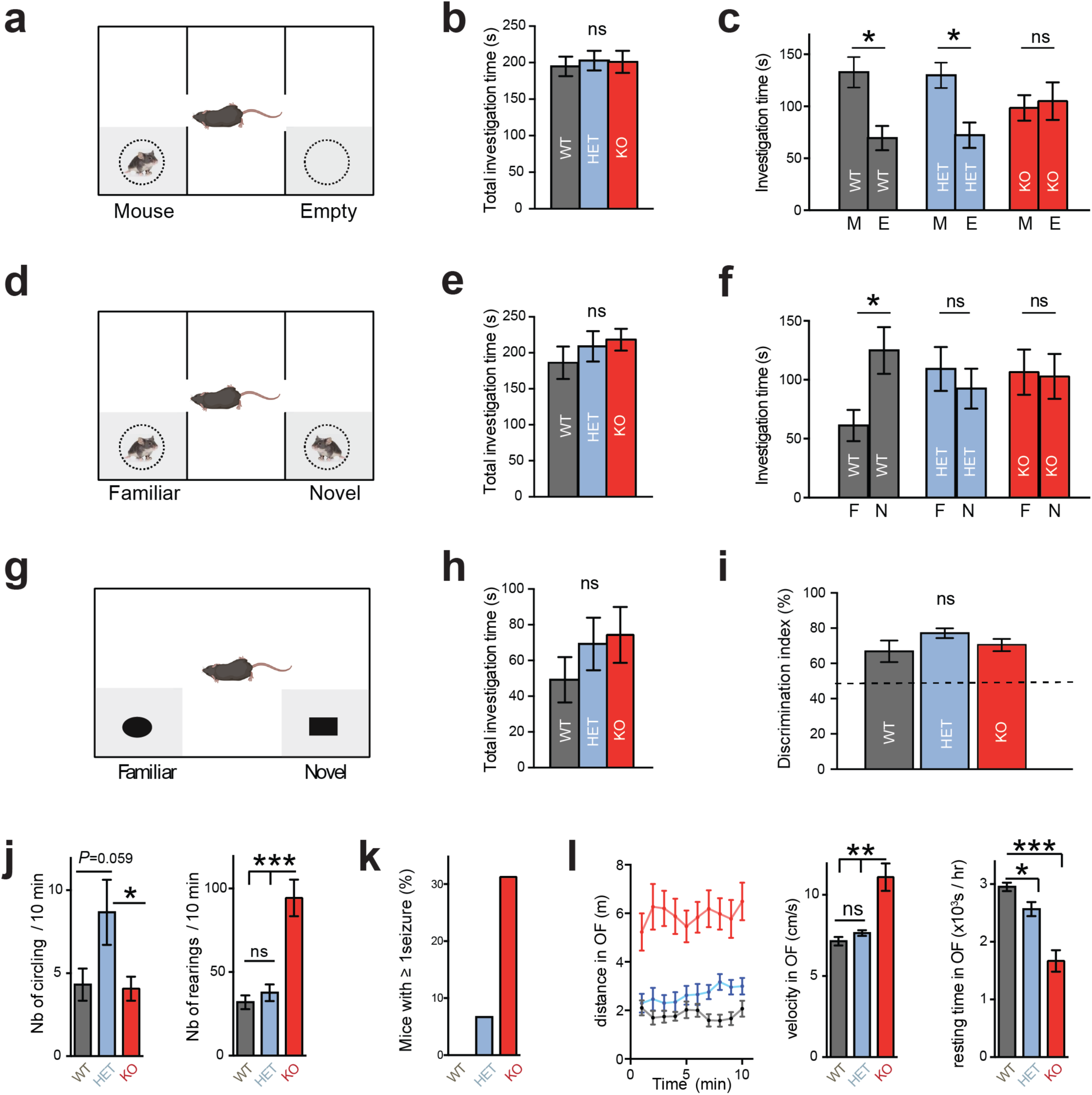
Behavioral hallmarks of neuropsychiatric disorders in *Neurod2* mutant mice. (**a-c**) During social interaction in the three-chamber test (**a**), all genotypes displayed similar total investigation time (**b**). However, while WT and HET mice spent around twice more time interacting with the mouse box than the empty box, *Neurod2* KO mice spent equivalent times investigating mouse and empty box (**c**). (**d-f**) Social recognition/memory test. When a familiar and a novel mouse were placed in each box (**d**), all genotypes spent similar times having social interactions (**e**), but only WT mice spent significantly more time investigating the novel mouse, indicating altered social recognition or memory in HET and KO mice (**f**). (**g-i**) In the novel object recognition test (**g**), *Neurod2* KO and HET mice displayed unaltered total investigation time (**h**) and discriminated normally between the familiar and the novel object (**i**). (**j**) Circling (left graph) and rearing (right graph) behaviors in the 3 genotypes. (**k**) Spontaneous seizures were observed in a third of *Neurod2* KO mice and in one HET mouse during the behavioral experiments, but never in WT littermates. (**l**) Hyperactivity in *Neurod2* KO mice. Left graph depicts the distance traveled in 1 minute-intervals during 10 minutes in the open field. Middle graph shows velocity during motion and right graph shows resting time, both during a 1-hour recording. N= 16 WT, 15 HET and 15 KO mice aged 8-14 weeks depending on the test. Data are means ± SEM. Statistical significance was evaluated by one-way ANOVA [(b), (e), (h), (i), (j), (k) and (l)] or two-way Mixed ANOVA followed by post hoc analysis using paired t-test [(c) and (f)] (ns, not significant; \**P* < 0.05; \*\**P* < 0.01; \*\*\**P* < 0.001).

**Figure 6:**
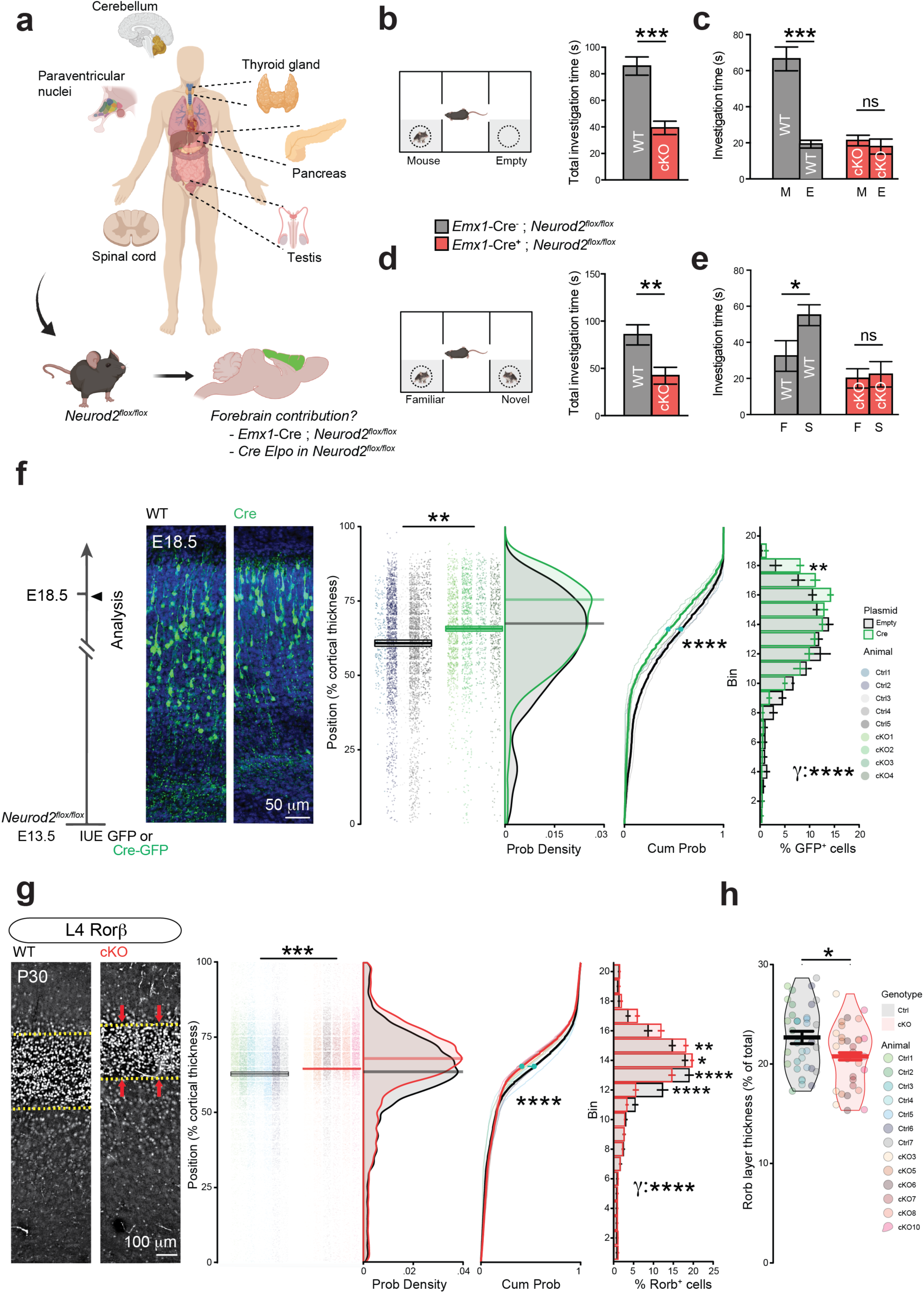
Forebrain excitatory neuron-specific *Neurod2* deletion recapitulates ASD-like phenotypes in mice. (**a**) Top scheme: Tissue types expressing *NEUROD2* in humans. They include paraventricular nuclei and thyroid hormone, which could account for ASD phenotypes in the *NEUROD2* syndrome. Hence, we generated *Neurod2*^flox/flox^ mice and crossed them with *Emx1*-Cre mice to assess the specific contribution of forebrain excitatory neurons to the ASD-related phenotypes (lower scheme). (**b-e**) Three-chamber assay. (**b,c**) Social interaction was altered. (**d,e**) Social recognition was reduced. Total interaction time is the time spent around left and right boxes. This parameter was altered, in contrast to the full KO mice. (**f**) Over-migration of CPNs is cell-autonomous, as shown by Cre electroporation in pallial progenitors at E13.5 followed by analysis at >E18.5. Mean cell laminar position in percentage of the cortical thickness, probability density function (bars representing maximal probability density), empirical cumulative distribution function and 20-bin based distribution of electroporated cells are represented from left to right. (**g**) Mean cell laminar position in percentage of the cortical thickness, probability density function (bars representing maximal probability density), empirical cumulative distribution function and 20-bin based distribution of CPN types using the markers Rorb for L4. (**h**) RORB layer thickness was decreased in *Emx1*-Cre; *Neurod2*^flox/flox^ mice. Data are means ± SEM. Statistical analyses were performed using two-tailed t-tests or Mann-Whitney test depending on the normality of samples [(b), (d), Y position in (f) and (g), (h)], by Kolmogorov Smirnov test [cumulative probability graphs in (f)-(g)] and by Permutation test for a spatially adjusted two-way ANOVA followed by Bonferroni’s post-hoc test [bin graphs in (f)-(g)] or by two-way Mixed ANOVA followed by post hoc analysis using paired t-test [(c) and (f)] (ns, not significant; \**P* < 0.05; \*\**P* < 0.01; \*\*\**P* < 0.001, \*\*\*\**P* < 0.0001).

Our study in the mouse, showing gene expression variation in the cerebral cortex, enrichment of ASD-related genes in orthologs of differentially expressed genes, functional alteration in neural circuits formed by CPNs and behavioral abnormalities associated with *Neurod2* conditional deficiency, provide some clues to the causative link between NEUROD2 deletions and ASD.

How neuronal migration and positioning contribute to ASD and by which transcription factors these biological processes are regulated is not well documented. Our data demonstrate that *Neurod2* cell-autonomously regulates radial migration and precise laminar positioning of CPNs. Cells migrated more superficially in *Neurod2* KO and *Emx1*-Cre; *Neurod2*^flox/flox^ mice than in corresponding control littermates. Whether *Neurod2* regulates multipolar-bipolar transition, multipolar neuron migration, glia-guided locomotion and/or terminal translocation remains to be investigated. Also, the target protein(s) downstream of *Neurod2* responsible for the migratory defect remain to be discovered. Similar to our study, excessive CPN migration has been reported after *in utero* overexpression of human *APP* (*45*), *Rac1* and its interactor *POSH* (*46*) or *Camk2b* (*47*). After *APP* overexpression however, over-migration was transient since it was not visible anymore at 6 days post-electroporation. This indicates that *APP* promotes migration velocity but does not regulate the final laminar positioning of CPNs. For *Rac1*, *POSH* and *Camk2b* however, the long-term impact of overexpression on cortical lamination was unexplored (*45–47*). Here, CPN over-migration in constitutive and forebrain excitatory neuron-specific KO mice was followed by an altered laminar positioning and thickness of layers, with mostly L5 expending superficially at the expense of L4. As a result, CPN subtypes displayed abnormally low *vs* high cell densities in deep *vs* superficial layers, respectively. This is to our knowledge the first time that such a phenotype is described. Given that even slight alterations in CPN positioning are sufficient to alter cortical output (*48*), this laminar positioning phenotype is expected to participate in disrupting cortical function. We observed only a slight corpus callosum size reduction and no obvious axonal targeting specificity defects in *Neurod2* null mice, contrasting with the total absence of corpus callosum in *Neurod2*/*NeuroD6* double mutant mice (*17*). A likely explanation resides in the functional redundancy between *NeuroD* family members concerning axon growth and navigation. In contrast, we suggest that the observed synaptic phenotypes are specific to *Neurod2* deletion among other *NeuroD* family members.

Our study indicates that some electrophysiological phenotypes are specific to CPN subtypes while other are generic. Although *Neurod2* is expressed in all CPN types, L2/3 CPNs in *Neurod2* KO mice display reduced frequencies of both mEPSCs and mIPSCs (*16*), while L5 CPNs show increased mIPSC amplitude (this study). Hence, *Neurod2* deletion appears to alter synaptic properties in a different manner depending on the CPN type and layer. In contrast, intrinsic excitability was increased similarly in L2/3 and L5 CPNs of *Neurod2* KO mice, suggesting that the control of excitability is a general function of *Neurod2*. This hypothesis is consistent with unpublished preliminary results indicating increased excitability of hippocampal CA1 PNs in *Neurod2* KO mice *in vivo* (Jérôme Epsztein, personal communication). However, the mechanisms by which *Neurod2* limits intrinsic excitability in L2/3 and L5 CPNs are likely different since decreased after-hyperpolarization was found as an underlying cause in L2/3 (*16*) but was unmodified in L5 CPNs (this study).

A literature-based search of our P30 differentially expressed gene list establishes a large number of *Neurod2* targets (direct or indirect) that might be causally associated with the cellular and behavioral phenotypes described in *Neurod2* KO mice.

Two particularly good candidates for the alteration of dendritic spine density and turnover in the apical tuft are the glucocorticoid receptor gene *Nr3c1* and the synaptome gene *Syne1* (CPG2 isoform), both downregulated in *Neurod2* KO mice. *Nr3c1* is a critical regulator of dendritic spine development and plasticity in CPN apical dendrites (*49*), a compartment where spine density was most affected in *Neurod2* KO mice. Acute and chronic stresses both preferentially alter spine plasticity in the apical tuft of CPNs through glucocorticoid binding to *Nr3c1* (*50*), which induces an internalization and transcriptional activity of *Nr3c1*. Interestingly, a recent study indicates that the transcriptional activity of *Nr3c1* requires *Neurod2* as a cofactor (*51*), and that *Neurod2* interacts even more strongly with the mineralocorticoid receptor *Nr3c2,* also involved in stress-dependent dendritic spine plasticity and a newly identified ASD factor (*41*). Together, these data suggest that *Neurod2* might be a nexus in stress-related synaptic plasticity in CPN apical dendrites. The hypothesis that *Neurod2* is involved in the stress pathway is reinforced by the fact that *Neurod2* KO mice show fearless behaviors in the elevated plus maze, fear conditioning (*8*) and conflict anxiety (this study), and by the ADHD symptoms found in 5/7 of the *NEUROD2*-mutated patients. Besides *Nr3c1*, the downregulated target *Syne1* isoform *CPG2* encodes an actin-binding protein regulating activity-dependent glutamate receptor endocytosis and RNA transport to nascent postsynaptic sites (*52*). Interestingly, pathogenic *CPG2* variants are associated with bipolar disorders (*53*). Although *Nr3c1* and *Syne1* are obvious candidates, other genes among the 31 post-synaptic and 4 pre-synaptic differentially expressed genes might participate in *Neurod2* spine phenotypes.

Concerning the increased CPN excitability, increased locomotor hyperactivity and propensity to spontaneous epilepsy observed in *Neurod2* KO mice that all suggest increased CPN activity, reasonable candidate genes lie among the 13 differentially expressed voltage-gated ion channels. *Kcnq5* and *Scn8a* are particularly good candidates, as both are restricted in CPNs and have been implicated in epilepsy and/or hyperactivity (*54, 55*).

Concerning increased Ih current density in L5 CPNs, a likely candidate gene from our RNA-seq is *Trip8b*. *Trip8b* is a critical regulator of membrane localization and expression of HCN1 channels that induce Ih current density (*56, 57*). Independently of its mechanism, increased Ih current density tend to decrease intrinsic neuronal excitability, suggesting that this phenotype is not part of initial defects but instead might represent a compensatory mechanism to reduce neuronal hyperexcitability (*27*).

The large number of *Neurod2* differentially expressed genes that can possibly account for the phenotypes observed in null mice renders functional rescue experiments impractical. In the future, since the brain alterations causing the phenotypes likely arise early in development it will be interesting to perform transcriptomic analyses at prenatal developmental stages. In recent years, there has been rapid advancement in single-cell transcriptomic technologies. Studying the transcriptomic changes in different CPN types and even in other cell types will provide valuable insights into the physiopathology of the *NeuroD2* syndrome.

The Emx1-Cre driver mouse line induces *Neurod2* deletion not only in the cortex but also in the hippocampus that expresses *Neurod2* at high levels. Hence, the contribution of the hippocampal formation to the behavioral defects in *Neurod2* deficient mice and *NEUROD2* mutated ASD patients remains to be investigated.

Finally, *in vitro* pieces of evidence have suggested a link between *Neurod2* and neuronal activity. Indeed, Neurod2 transactivation is increased by activity in cultured cortical neurons (*10*), and *Neurod2* mRNA expression is regulated by NMDAR activation (*58*). In this context, it will be interesting to test whether and how *Neurod2* regulates experience-dependent neuronal and synaptic development *in vivo*, and how this relates to neuropsychiatric disorders.

This study, from human to rodent model, identifies *NEUROD2*/*Neurod2* as a new gene linked to ASD, essential for CPN development and function. Indeed, its deletion affects radial migration, the cortical expression of a number of genes related to ASD and induces ASD-relevant deficits, associated with layer positioning and thickness defects, structural changes at synapses formed by deep-layer CPNs and functional changes in excitability without obvious alterations in neuron viability and path-finding. Our data point to NEUROD2 as a key member of a transcriptional regulatory network whose alteration can lead to a convergence of brain phenotypes centering on ASD and to murine *Neurod2* mutants as new candidate animal models of ASD.

## Acknowledgements

This work was supported by the Agence Nationale de la Recherche (ANR-13-JSV4-0006 to A.d.C. and ANR-13-BSV4-0013-01 to H.C.), the Fondation Pour la Recherche sur le Cerveau and the Fondation Lejeune (both to A.d.C.), the European Community 7th Framework programs Development and Epilepsy—Strategies for Innovative Research to improve diagnosis, prevention and treatment in children with difficult to treat Epilepsy [DESIRE] (to A.d.C., C.C and A.R.) and the Fritz Thyssen Stiftung Foundation (10.15.2.022MN to C.C.). High throughput sequencing was performed at the TGML Platform, supported by grants from Inserm, GIS IBiSA, Aix-Marseille Université, and ANR-10-INBS-0009-10. Financial support to S.B. was provided by the Fondation Pour la Recherche Médicale. We thank the PPGI, PBMC and InMAGIC platforms at INMED for their important technical support, J.B. Manent, O.J. Manzoni and P. Chavis for advice along the course of the study, Bruno Pichon and Audrey Van Hecke for assistance in genetic and clinical data, respectively. The project that gave rise to these results received the support of a fellowship from “la Caixa” Foundation (ID 100010434). The fellowship code is LCF/BQ/PI19/11690010.

## Author contributions

A.d.C. and H.C. conceived and initiated the project. K.R., R.M. S.B. and A.d.C. conceived and designed all the experiments. A.R., H.C. and C.C. provided critical input to the project at all steps. K.R., R.M., S.B., S.L., C.B., S.S., F.S., A.L., L.H., S.G., M.C., E.P.P., A.M. and A.d.C. performed the experiments. Lamination experiments were done by R.M. and K.R.; migration experiments by K.R. and F.S., spine density experiments by S.L. and A.d.C., spine turnover experiments by S.B., electrophysiological synaptic properties experiments by S.S., intrinsic neuronal properties by C.B., RNAseq experiments by S.L., L.H. and A.d.C., behavior by S.G. (Phenotype Expertise), qPCR and Sanger sequencing by E.P.P. and A.M. and *in vitro* pathogenicity testing experiments by E.P.P., R.M. and A.d.C. All analyzes were performed by K.R., R.M., S.B. or A.d.C. Scientific discussions and collaborative experiments with A.B. grounded the project. J.A.R., E.H., K.L., S.M.A., L.J., A.V.H., O.V., B.P., A.V.H., D.A., S.K., and C.G. recruited patients, acquired clinical data, or analyzed whole-exome or Sanger sequencing results. K.R., R.M., S.B. and A.d.C. drafted the original manuscript, and all authors assisted in editing the manuscript.

---

**Supplementary Figure 1:**
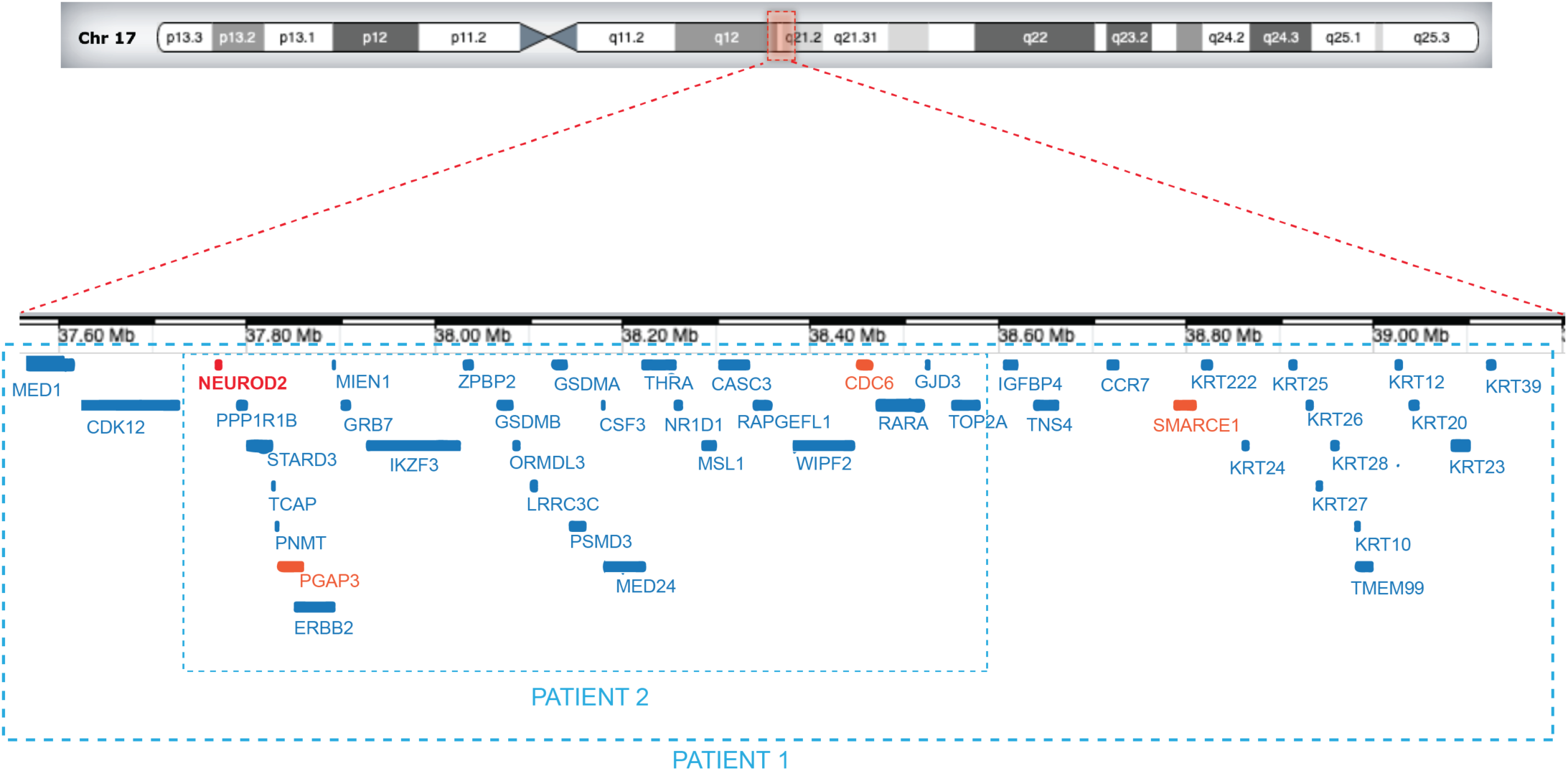
Schematic representation of 17q12 deletions in patients 1 and 2 (complement to Figure 1)

**Supplementary Figure 2:**
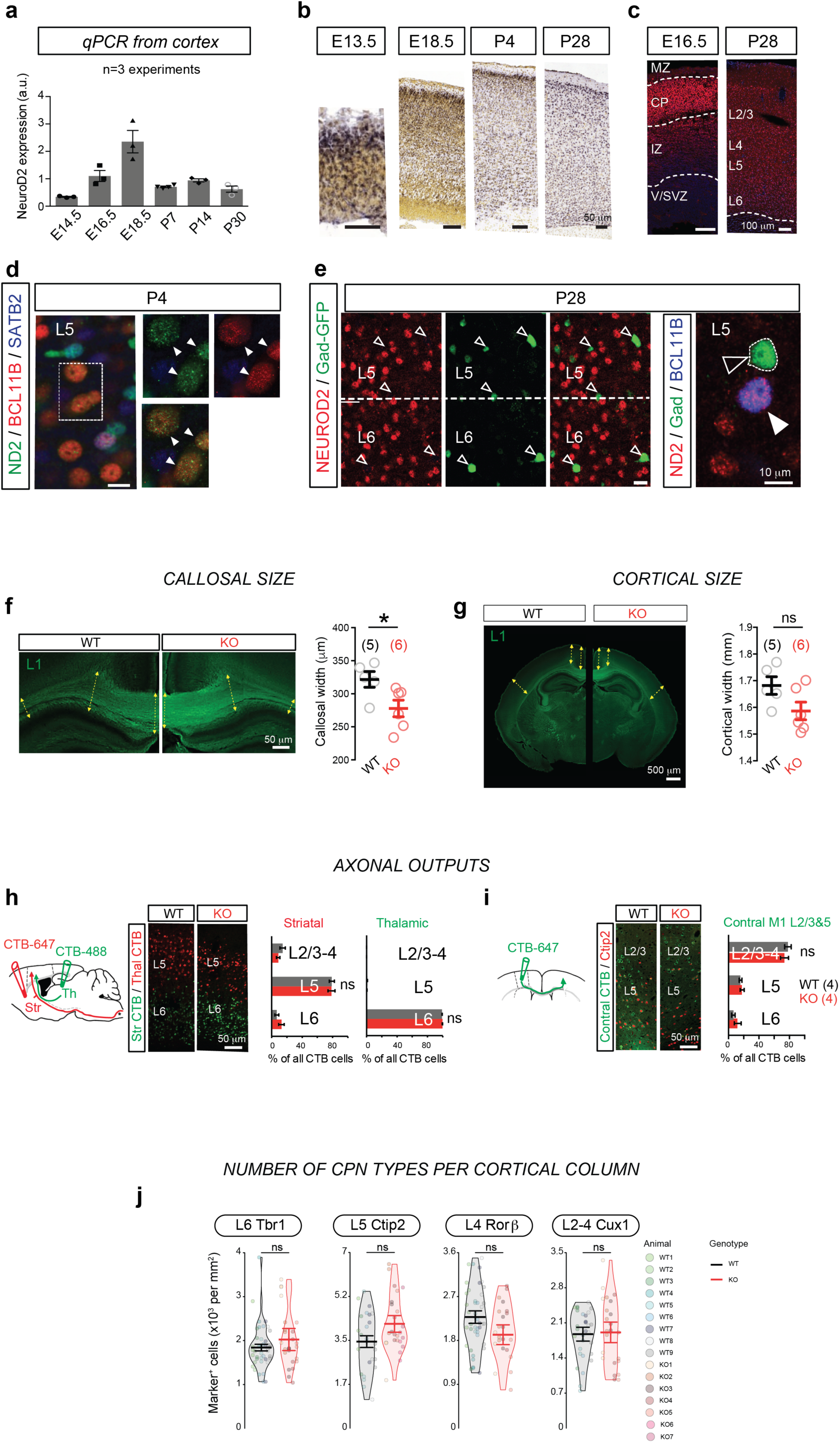
Developmental expression, axonal projection and number of CPN types in *Neurod2* KO mice (complement to Figure 2). (a-e) Cortical expression of NeuroD2 across development. (a) qPCR shows permanent cortical expression of *Neurod2* mRNA with a peak expression at E18.5. (b) In situ hybridizations from Allen Brain Atlas confirm permanent *Neurod2* mRNA expression across cortical layers and ages (S1 shown). (c) NEUROD2 protein expression at E18.5 and P28 in S1. (d) Neurod2 protein (green) is co-expressed with BCL11B (red) at P4 in L5. (e) At P28, Neurod2 (red) never co-localizes with Gad67-GFP (green) in L5 and L6 (f), but co-localizes with BCL11B in L5 (g). (f-i) Callosal size and cortical size. Callosal width was slightly decreased (f) but cortical width was unaltered (g) in *Neurod2* KO mice. (h,i) Axonal targeting specificity of CPN subtypes. Fluorescent retrograde cholera toxin beta injection in striatum, thalamus (h) and contralateral M1 L2-5 (i) led to unaltered distribution of retrogradely-labeled somata in ipsilateral motor/somatosensory cortex. (j) Quantity of CPN subtypes per cortical column was not affected, indicating normal production and unaltered apoptosis. Data are represented as means ± SEM. Statistical significance was evaluated by Student t-test or Mann-Whitney test depending on normality of samples [(f), and (j)] or by two-way ANOVA followed by Bonferroni’s post-hoc test [(h), (i)] (\**P* < 0.05).

**Supplementary Figure 3:**
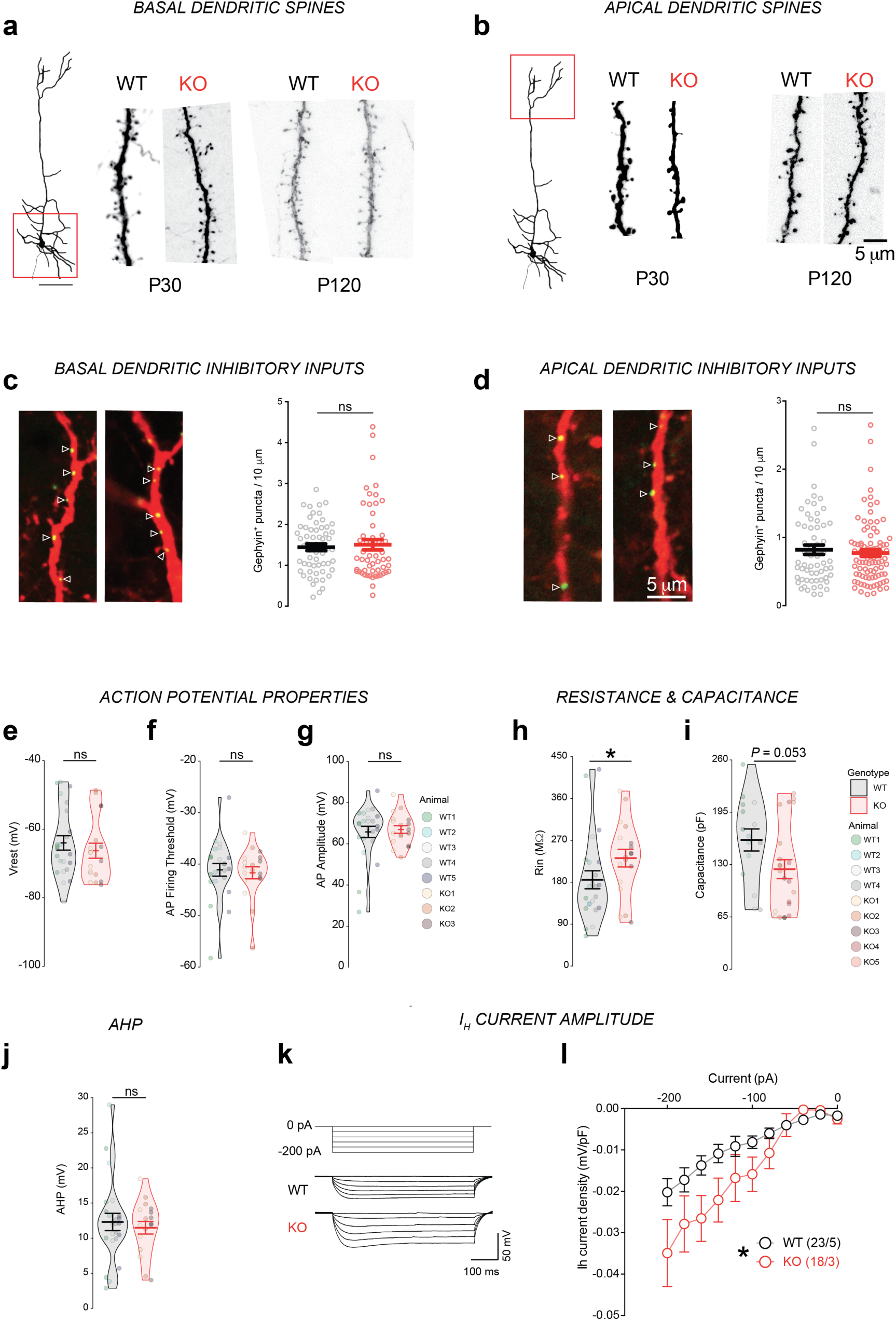
Synaptic and physiological defects in CPNs of *Neurod2* KO mice (complement to Figure 3) (a,b) Confocal images of dendrites of basal (a) and apical (b) dendritic spines of *Neurod2* WT and KO mice at P30 and P120 (complement to Figure 3a-c). (c-j) Intrinsic properties of L5 CPNs in *Neurod2* KO mice (complement to Figure 3i-m). (c-e) Action potential properties, including Vrest (c), AP firing threshold (d) and AP amplitude (e). (f,g) Membrane resistance (f) and capacitance (g). (h) After-hyperpolarization (AHP) was normal in L5 PNs of *Neurod2* KO mice. (I,j) *Neurod2* KO L5 PNs exhibited increased Ih-current amplitudes compared with WT PNs (current/voltage relation of Ih currents). (i) shows experimental protocol and representative traces, (j) shows summary graph of the voltage-current relation. Data are means ± SEM. Statistical analyses were performed using two-tailed t-tests or Mann-Whitney test depending on the normality of samples [(c) to (g)] and by two-way repeated measure ANOVA followed by Bonferroni’s post hoc test (j). \**P* < 0.05.

**Supplementary Figure 4:**
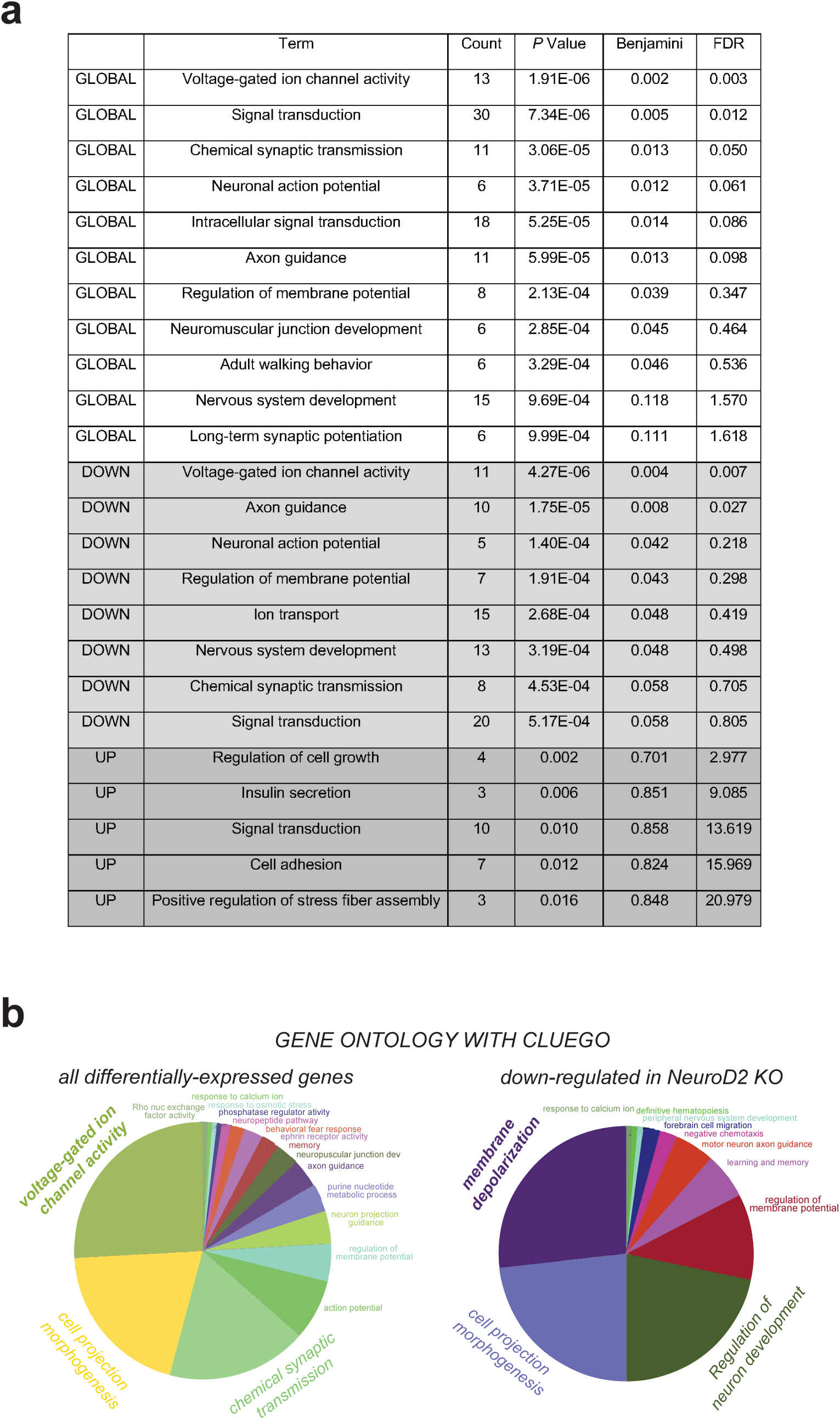
Gene ontology (complement to Figure 4) (a) Gene ontology of differentially expressed genes in M1/S1 of *Neurod2* KO mice with DAVID (https://david.ncifcrf.gov/). GLOBAL: all differentially expressed genes. DOWN: down-regulated genes. UP: up-regulated genes. FDR: False Discovery Rate. (b) GO pie charts from ClueGO (Cytoscape).

**Supplementary Figure 5:**
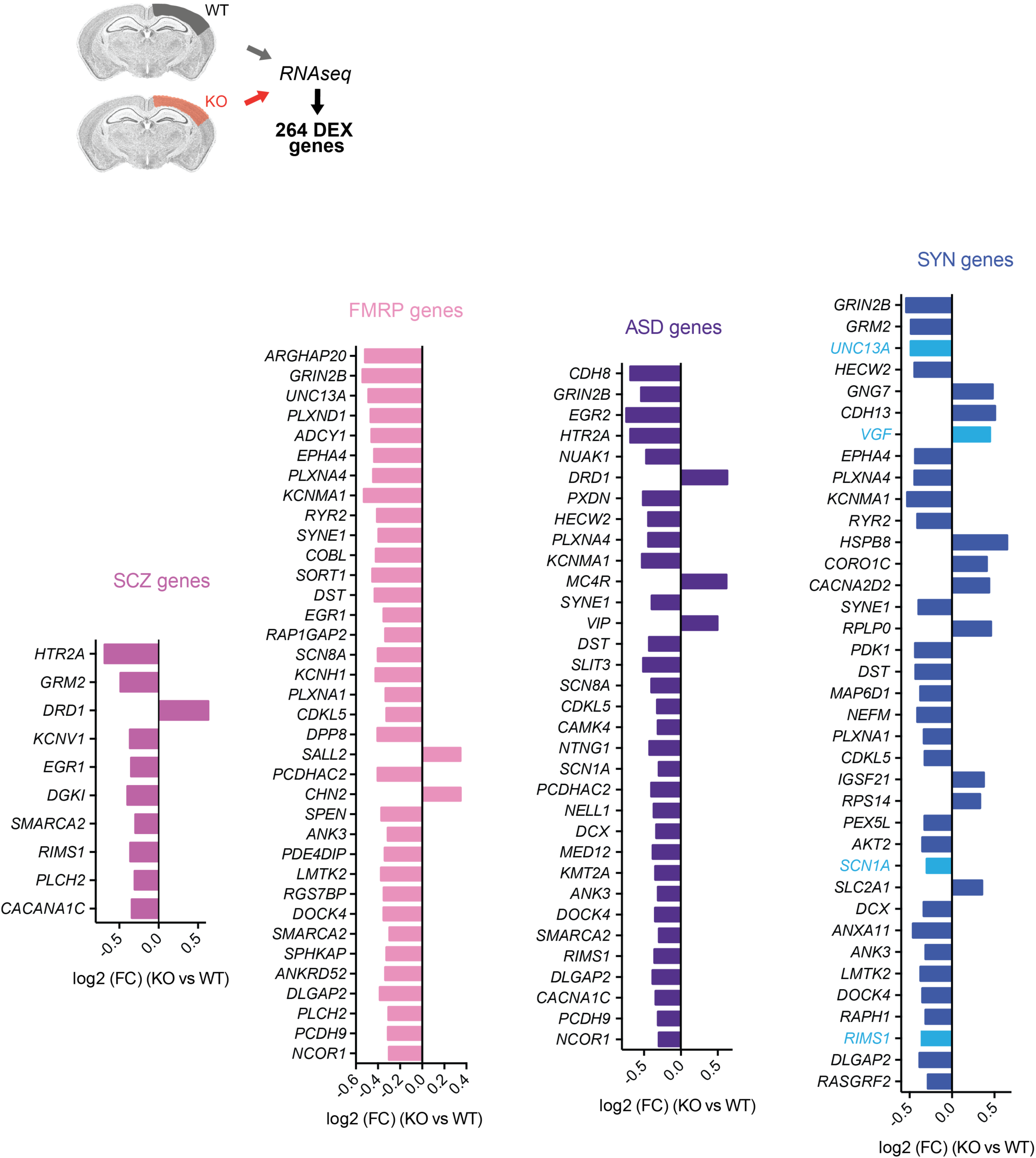
*Neurod2* KO differentially expressed genes associated with neuropsychiatric disorders (complement to Figure 4) Fold change expression (FC; log2 scale) of differentially expressed genes belonging to specific gene sets, ranked according to adjusted *P* value (lowest *P*adj on top). In the graph representing synaptome genes (SYN), presynaptic genes are depicted in light blue while postsynaptic genes are in dark blue.

**Supplementary Figure 6:**
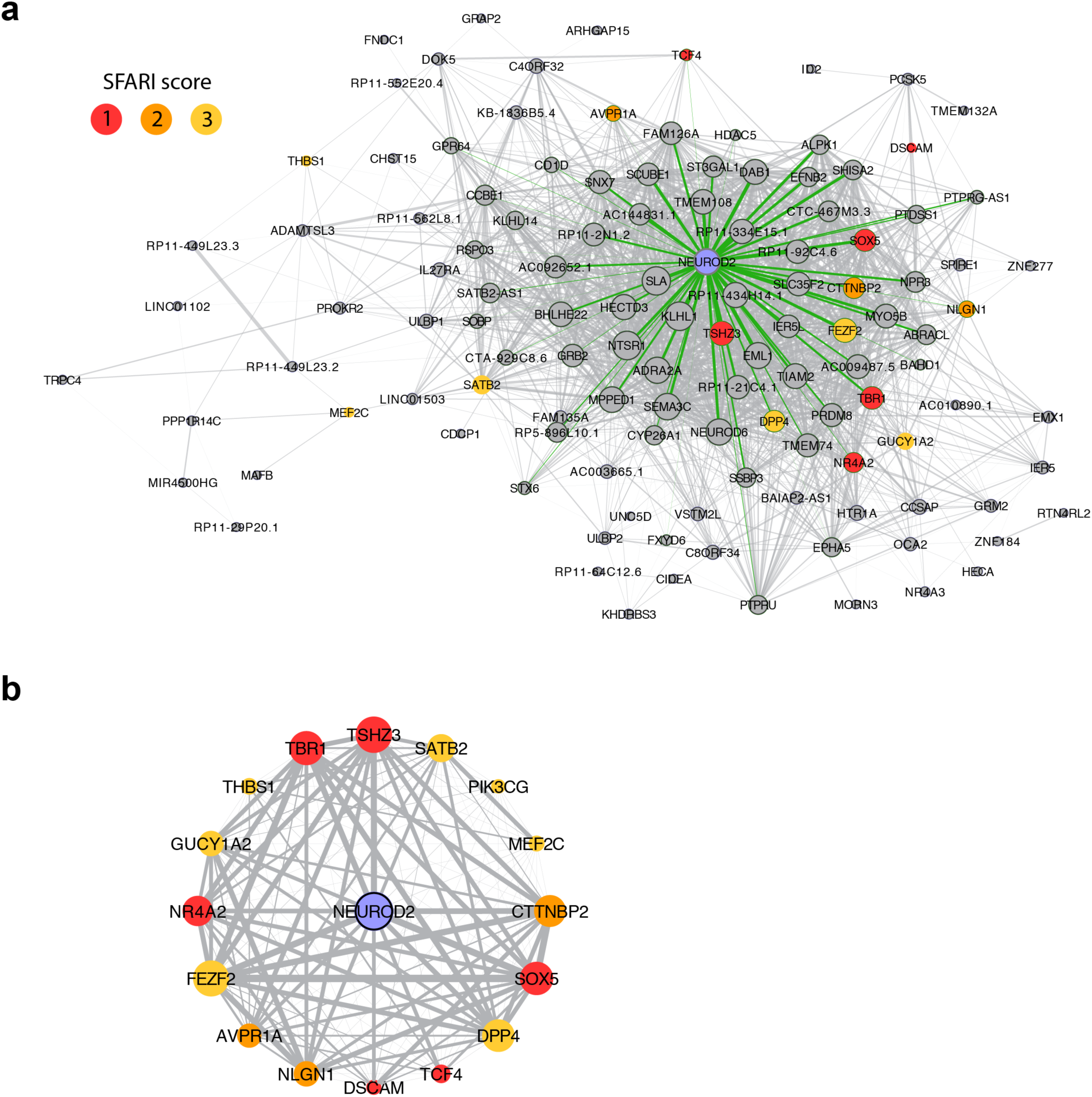
NEUROD2-based coexpression network analysis in humans (complement to Figure 4). (a) Network representation of NEUROD2 within module 37 from Li et al. 2018 [ref 51] showing gene connectivity based on Pearson correlation (weight of edges scaled for r between 0.7 and 1). The size of each node is proportional to degree centrality calculated with a cutoff of r=0.7. Green edges indicate direct neighbors of NEUROD2 with threshold r=0.7. Genes described in SFARI are colored according from SFARI score (1 to 3). (b) Circular network representation of NEUROD2 and SFARI genes interactions with weight of edges scaled for r between 0.5 and 1. Size of the node represent the same values of degree centrality of calculated in (a).

**Supplementary Figure 7:**
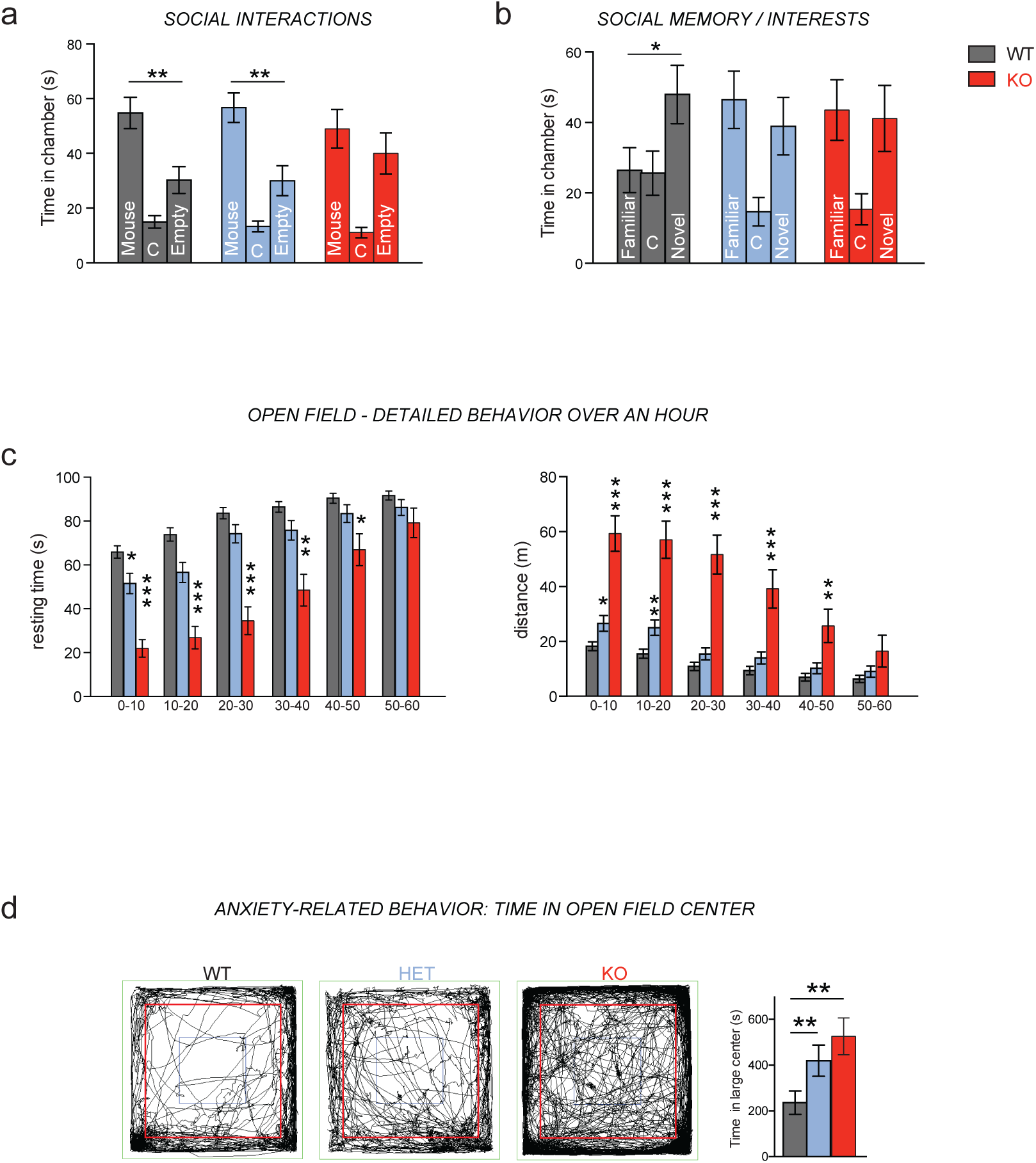
Behavioral phenotypes in *Neurod2* KO and HET mice (complement to Figure 5). (a,b) Time spent in the different chambers during the 10 minutes social interaction (a) and social memory tests. (c) Resting time and distance during 10 minutes intervals over the 1-hour open field assessment. (d) Time spent in the large center of the open field (red square in representative traces) was significantly increased in both HET and KO mice. Data are means ± SEM. Statistical significance was evaluated by one-way ANOVA (d) or two-way Mixed ANOVA followed by post hoc analysis using paired t-test [(a) to (c)] (ns, not significant; \**P* < 0.05; \*\**P* < 0.01; \*\*\**P* < 0.001).

**Supplementary Figure 8:**
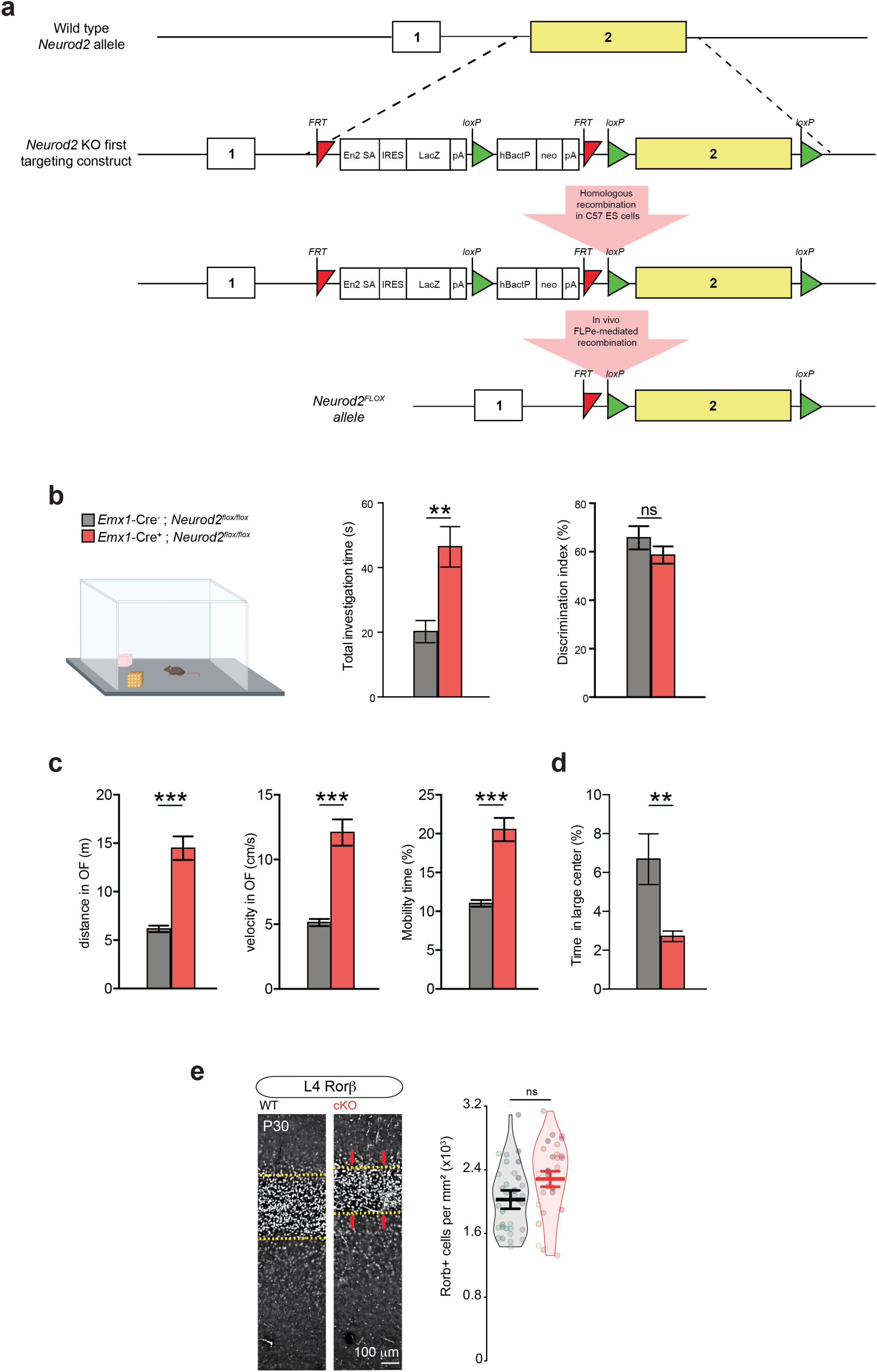
Forebrain excitatory neuron-specific *Neurod2* deletion recapitulates ASD-like phenotypes in mice (complement to Figure 6) (a) Schematic of strategy to generate C57BL/6 mice carrying the Neurod2 FLOX allele. (b) *Emx1*-Cre; *Neurod2*^flox/flox^ mice displayed increased total investigation time and discriminated normally between the familiar and the novel object. (c) Hyperactivity in *Emx1*-Cre; *Neurod2*^flox/flox^ mice. Left graph depicts the distance traveled in 20 minutes in the open field. Middle graph shows velocity during motion and right graph shows the percentage of time during which the mouse is mobile. (d) Time in large center (%) is decreased in *Emx1*-Cre; *Neurod2*^flox/flox^ mice. (e) Density of RORB-expressing cells in *Emx1*-Cre; *Neurod2*^flox/flox^ *vs* control littermates. N= 10 mice aged 8-14 weeks depending on the test. Data are means ± SEM. Statistical analyses were performed using two-tailed t-tests or Mann-Whitney test depending on the normality of samples (ns, not significant; \**P* < 0.05; \*\**P* < 0.01; \*\*\**P* < 0.001).

